# Quinolinic acid phosphoribosyl transferase moonlights as an apoptosis regulator to empower lung cancer progression

**DOI:** 10.64898/2026.04.01.715697

**Authors:** Hossein Kashfi, Didem Ilter, Angelo A. Nicolaci, John Lockhart, Stanislav Drapela, Felicia Lazure, Devesh Raizada, Nadir Sarigul, Julia D. Spegel, Nathan P. Ward, Tumpa Dutta, Stephen J. Gardell, Jennifer M. Binning, Elsa R. Flores, Gina M. DeNicola, Ana P. Gomes

**Affiliations:** Department of Tumor Microenvironment and Metastasis, H. Lee Moffitt Cancer Center & Research Institute, Tampa, FL, USA; Department of Molecular Oncology, H. Lee Moffitt Cancer Center & Research Institute, Tampa, FL, USA; Department of Metabolism and Physiology, H. Lee Moffitt Cancer Center & Research Institute, Tampa, FL, USA; Translational Research Institute, Advent Health, Orlando, FL 32804, USA

## Abstract

Although nicotinamide adenine dinucleotide (NAD⁺) metabolism is fundamental for cancer cell survival, the role of the *de novo* NAD⁺ biosynthetic pathway, particularly in non-small cell lung cancer (NSCLC), remains largely unknown. Here, we describe a non-canonical role for the rate-limiting enzyme in *de novo* NAD^+^ biosynthesis, quinolinate phosphoribosyltransferase (QPRT), in NSCLC progression. We show that QPRT is highly expressed in late-stage tumors and required for NSCLC growth; however, its suppression does not change NAD⁺ levels or elicit compensatory NAD⁺ biosynthetic activity. Instead, QPRT interacts with caspase-3 and suppresses its activation, protecting NSCLC cells from apoptosis. This reveals a moonlighting function for QPRT in apoptosis regulation independent of its enzymatic activity in tryptophan catabolism. Together, these findings, redefine QPRT as a protein with dual functionality and reveal it as a potential therapeutic target in NSCLC, highlighting the importance of non-canonical roles of metabolic enzymes in cancer biology.

**Significance:** This study reveals that QPRT supports NSCLC progression by directly inhibiting caspase-3–mediated apoptosis independent of NAD biosynthesis, redefining its role and highlighting non-enzymatic functions of metabolic enzymes as therapeutic targets.

## Introduction

Lung cancer is still the leading cause of cancer-related mortality worldwide (1). Non-small cell lung cancer (NSCLC) accounts for approximately 85% of lung cancer cases (2). In recent years there has been many advances in targeted and immune-based therapies yet most patients eventually relapse, highlighting the urgent need for more effective therapeutic strategies (2). Metabolic reprogramming in cancer cells is a hallmark of cancer, and is recognized as a driver of tumorigenesis and therapy resistance (3). Altered NAD⁺ metabolism is one of these metabolic adaptations that is essential in supporting cancer cell survival, redox homeostasis, and adaptive stress responses, and thereby presents itself as a potential therapeutic vulnerability in NSCLC (4–6). NAD^+^ not only acts as an electron carrier in redox reactions but also functions as a co-substrate for regulatory enzymes that regulate genomic stability, stress resistance and metabolic adaptation (5,7). NAD⁺ has an indispensable role in maintaining cellular energetics since its reduced form, NADH, donates electrons to the electron transport chain and fuels mitochondrial respiration (8). Recent studies have demonstrated that NAD⁺ levels play a critical role in NSCLC cells, enabling them to sustain high metabolic demand and resist therapeutic stress (9–11). To meet their NAD⁺ demands mammalian cells rely on three major biosynthetic pathways: the *de novo* synthesis pathway that fuels NAD^+^ from tryptophan (Trp), the Preiss–Handler pathway that uses nicotinic acid (NA) as the NAD^+^ precursor, and the salvage pathway, which recycles dietary and endogenous precursors such as nicotinamide (NAM) (12).

Most mammalian cells, including cancer cells, are thought to meet their NAD^+^ demands through the salvage pathway (7). In line with this, nicotinamide phosphoribosyl transferase (NAMPT), the rate-limiting enzyme in the NAD⁺ salvage pathway, is frequently overexpressed in various human cancers, including NSCLC, and is often associated with poor prognosis (13–15). Thus, pharmacological inhibition of NAMPT has been put forward as a strategy to treat cancer (15,16). However, despite remarkable efficacy in driving cancer cell death *in vitro*, NAMPT inhibitors have failed to produce an effective therapeutic response *in vivo* (17), suggesting that alternative pathways can fuel NAD^+^ levels in cancer cells in the absence of the salvage pathway (6). While the NAD^+^ salvage pathway has been extensively studied in the context of cancer(18–21), including NSCLC(9,22,23), the role of *de novo* NAD^+^ biosynthesis is understudied. In the *de novo* NAD^+^ biosynthetic pathway, Trp is converted through a multi-step process into quinolinic acid (QA), which is then metabolized by quinolinic acid phosphoribosyl transferase (QPRT) into nicotinic acid mononucleotide (NAMN), a key intermediate in NAD⁺ biosynthesis. Thus, QPRT serves as the rate-limiting enzyme that bridges Trp catabolism to NAD⁺ production through the *de novo* route (24,25). While this pathway is primarily active in the liver, its contribution to systemic NAD⁺ levels vary by tissue and is relatively modest in the heart and other peripheral organs (12,26). Although QPRT plays a key role in physiology by regulating the rate of *de novo* NAD^+^ biosynthesis (12,26), its specific contributions to NSCLC metabolism and consequent effects on tumor progression remain underexplored. Here, we aimed to elucidate the functional significance of QPRT in NSCLC and its potential as a therapeutic target.

## Materials and Methods

### Experimental mice

All animal experiments were approved by the Institutional Animal Care and Use Committee (IACUC) of Moffitt Cancer Center managed by the University of South Florida. Animals were maintained in an AAALA-accredited animal facility at 20–23 °C with 30–70% humidity on a 12 h light-dark cycle. To generate animals with lung adenocarcinoma tumors, Kras^LSL-G12D/+^; Trp53^flox/flox^ (KP) (27) and Kras^LSL-G12D/+^; Lkb1^flox/flox^ (KL) (28) mice were infected intranasally with Ad5CMVCre or Ad5mSPC-Cre adenovirus (University of Iowa, Iowa City, IA), respectively. For xenograft experiments with human cells immunodeficient NOD *scid* gamma (NSG) mice, bred and maintained in the Gomes lab colony, were utilized.

### Cell lines and cell culture

PC9, H23, H441 H1581, H1648, H1650, H2009, H2030, H2087, H2122, and H2228 NSCLC cell lines were obtained from the Hamon Cancer Center Collection (University of Texas-Southwestern Medical Center) or the Lung Cancer Center of Excellence at Moffitt Cancer Center, or purchased from ATCC. NSCLC cells, unless mentioned otherwise, were maintained in RPMI 1640 medium (Cytiva, SH30027FS) supplemented with 5% fetal bovine serum (FBS; Gibco, A52567-01) and 1% penicillin-streptomycin (Cytiva, SV30010), at 37°C in a humidified incubator with 5% CO₂. H2122 and H2228 were cultured in RPMI 1640 supplemented with 10% FBS and 1% penicillin-streptomycin. KP 1.9 and KL 95 cell lines were obtained from Dr. Alfred Zippelius (University of Basel) and HEK293T cells were bought from GenHunter, all three cell lines were maintained in high glucose DMEM media with L-glutamine (Cytiva, SH30022) supplemented with 10 % FBS (FBS; Gibco, A52567-01) and 1% penicillin/streptomycin (Cytiva, SV30010).

### shRNA mediated knockdown

pLKO.1 based shRNA constructs (non-targeting shNT: SHC002, shQPRT #1: TRCN0000034846, shQPRT #2: TRCN0000034848) were obtained from Sigma. pMD2.G (Addgene plasmid 12259) and psPAX2 (Addgene plasmid 12260) constructs were co-transfected with each shRNA construct into HEK293T cells growing in their standard media as mentioned above using the X-tremeGENE (Roche) reagent according to the manufacturer’s protocol. The media was refreshed 24 h after transfection and media containing lentivirus was collected 72 h after the media change. The collected media containing viral particles was filtered using 0.45 µm syringe filters and used for transducing human cell lines in the presence of 10 µg/mL polybrene. NSCLC cells were selected with 2 µg/mL puromycin starting 24 h after transduction. Cells were maintained with puromycin for the duration of the experiments to prevent drift.

### Cell proliferation and cytotoxicity assays

For knockdown experiments, cells were transduced overnight with lentiviruses packaged with either non-targeting shRNA or shRNA against QPRT constructs. The following day, selection was started with puromycin (2 μg/mL). On day 4 post-transduction, when selection process is confirmed to be completed on a non-transduced kill plate, cells were seeded into 96-well plates for growth and cytotoxicity assays for three days at the following densities: H2009 and PC9 at 6,000 cells/well, H2030 at 2,500 cells/well, and H1581 at 12,000 cells/well.

Cell proliferation was monitored using the Incucyte live-cell imaging system. 3 days after seeding, cytotoxicity was assessed at endpoint using either 2.5 μg/mL propidium iodide (PI) (Thomas Scientific, C974V80) or 50 nM SYTOX Green 50 nM (Thermo Scientific, S7020) as indicated in figure legends. For the apoptosis assay, a similar scheme as described above was followed for cell seeding but cells were seeded and maintained in the presence of 0.25 µg/mL Annexin V fluorescent conjugates (CF450; Biotium, 29083) for the following 3 days. In these experiments, cell death was measured as percentage of PI^+^ or SYTOX Green^+^ stained cells over total number of cells determined by the cell-by-cell analysis by the Incucyte Software (v2021A). Similarly, for apoptosis analysis percentage of Annexin V-stained apoptotic cells over total number of cells were calculated.

### 3D spheroid culture and cytotoxicity assay

QPRT knockdown was achieved as described above. Four days following transduction, PC9 (200 cells/well) and H1581 (2,000 cells/well) were seeded in ultra-low attachment 96-well plates (Fisher Scientific, 7007) to establish 3D spheroid cultures. Spheroid formation and growth were monitored over 4–6 days using the Incucyte live-cell imaging & analysis system. Cytotoxicity was assessed with SYTOX Green (50 nM) staining, an indicator of cell death, and measuring percentage of SYTOX Green positive area of the total spheroid area. The level of apoptosis was assessed similarly by staining with 0.25 µg/mL Annexin V CF450 fluorescent conjugates (Biotium, 29083) instead.

### Flow cytometry

The knockdown of QPRT in NSCLC cells was performed as previously described in above section.. For DiD labeling and proliferation assay, four days after transduction, cells were stained with DiD dye (Thermo Fisher Scientific, D7757) in PBS without calcium or magnesium. H2030 and H2009 cells were stained with 5 µM DiD, and PC9 cells with 3 µM DiD, for 30 minutes at 37°C. Following staining, cells were trypsinized, pelleted by centrifugation (300g x 5 min), and approximately 100,000 cells were collected for baseline DiD median fluorescence intensity measurement (day 0) using flow cytometry (Attune NxT, Thermo Fisher). The remaining cells were reseeded for the experiment. Four days later, cells were harvested and analyzed again by flow cytometry using the same procedure. Proliferation was assessed by quantifying dye dilution over time, normalized to day 0 fluorescence intensity.

Propidium iodide (1 µg/mL) was included to discriminate dead cells. For *in vitro* cell cycle analysis, Vindelov assay was performed. Cells were trypsinized, washed with PBS, resuspended in 0.5 mL PBS, and then fixed by slowly adding 4.5 mL of 70% ethanol. The Vindelov staining solution contained 10 mM Tris buffer (Research Products International, T60040-10000.0), 20 µg/mL RNase A (Millipore Sigma, 10109142001), 0.6 mg/mL NaCl (Sigma-Aldrich, S3014-5KG), and 50 µg/mL PI prepared in distilled water. Following 24 hours of fixation at 4°C, cells were centrifuged (500 x *g* x 5 min), washed twice to remove residual ethanol, resuspended in the Vindelov solution, and incubated for 1 hour at 37°C. Fluorescence was then recorded on a BD Canto cytometer with a low event acquisition rate.

### Total NAD^+^ measurements

On day one post transduction, selection for cells with QPRT knockdown was started with puromycin (2 µg/mL) and seeded in 96-well plates at the following densities: 8,000 cells/well for PC9 and H2009, 5,000 cells/well for H2030, and 14,000 cells/well for H1581. 48 hours after seeding, total NAD^+^ levels were quantified using the NAD/NADH-Glo Assay (Promega, G9072) according to the manufacturer’s instructions. Cells were plated in a separate plate under the same conditions to be able to measure DNA content using CyQUANT Cell Proliferation Assay kit according to the manufacturer’s protocol (Thermo Fisher, C7026) for normalizing NAD^+^ levels.

To determine if an NAD^+^ precursor influences the amount of NAD^+^ in cells with QPRT knockdown, NSCLC cells were treated with nicotinamide riboside (NR, 1 mM in DMSO) (MedChem Express, HY-123033A) or β-Nicotinamide mononucleotide (NMN), 1 mM in water, MedChem Express, (HY-F0004) or vehicle (DMSO/water) 6 hours after they were transduced with lentiviruses to express either shQPRT or non-targeting control. The next day, cells were subjected to puromycin selection and seeded for the assay while being treated with either NR, NMN or vehicle (DMSO/water) with the following seeding densities: PC9 and H2009 cells 8000 cells/well, and H2030 5000 cells/well of 96-well plates. The cells were then collected for the NAD assay 72 h after transduction. NAD^+^ levels were normalized using DNA context as described above.

To determine whether NAD⁺ levels were altered under tryptophan-restricted conditions, one day after transduction, H2030 cells were selected with puromycin and seeded in 96 well plate (5000 cells/well). The next day the medium was changed to RPMI 1640 medium lacking L-tryptophan and supplemented with dialyzed fetal bovine serum (dFBS; Sigma-Aldrich, F0392) to remove residual amino acids. For restricted conditions, L-tryptophan was added back at a final concentration of 0 mg/L and 1.25 mg/L compared to normal levels of 5 mg/L in standard RPMI 1640. The cells were harvested 72 hours after transduction for NAD^+^ measurement as described above.

To assess if NAD^+^ precursors can rescue the reduction in NAD^+^ levels upon disruption of the salvage pathway with the NAMPT inhibitor FK866, parental PC9, H2009 and H2030 cells were seeded in 96-well plates at the following densities: H2009 and PC9 at 5,000 cells/well, H2030 at 1,500 cells/well, and H1581 at 10,000 cells/well. The next day, cells were treated for 72 hours with NAD^+^ precursors, nicotinic acid (NA, 20 µM; Sigma-Aldrich, N0761), quinolinic acid (QA, 20–100 µM; Sigma-Aldrich, P63204), nicotinamide riboside (NR, 1 mM; MedChem Express, HY-123033A) and L-tryptophan (Trp, 32 µM; Sigma-Aldrich, T0254) in the presence or absence of FK866 (20 nM; Cayman Chemical Company, 13287). Normalized NAD levels were measured as described above.

### Z-VAD-FMK and Ferrostatin-1 treatment for cell death inhibition

NSCLC cells with QPRT knockdown were generated as described above. On day 3 post-transduction, cells were seeded into 96-well plates at the following densities: H2009 and PC9 at 5,000 cells/well, H2030 at 1,500 cells/well, and H1581 at 10,000 cells/well. The following day cells were treated with Z-VAD-FMK (50 µM for PC9, H2009 and H2030 and 100 µM for H1581) (Fisher Scientific, 50-112-6908), a pan-caspase apoptosis inhibitor, or Ferrostatin-1 (20 µM) (Cayman Chemical Company, 17729), a ferroptosis inhibitor, or vehicle (DMSO) for 72 hours. After the treatment period, cell death was assessed as described above using PI staining.

### Caspase 3/7 activity measurement

QPRT knockdown cells were generated as described earlier. Four days after transduction, cells were seeded in the presence of puromycin in 96-well plates at the following densities: H2009 and PC9 at 6,000 cells/well, H2030 at 2,500 cells/well, and H1581 at 14,000 cells/well. Cells were incubated for 72 h before measuring caspase-3/7 activity according to the manufacturer’s protocol (Promega, G8091). The measured activity was normalized to the DNA content using CyQUANT Cell Proliferation Assay kit according to the manufacturer’s protocol (Thermo Fisher, C7026).

To assess caspase-3/7 activity following treatment with NAD^+^ precursors NR (1 mM) or NMN (1 mM), QPRT knockdown cells were generated in the presence of NR (1 mM) or NMN (1 mM) as described above. Four days after transduction, cells were seeded into 96-well plates at the following densities: PC9 and H2009 at 6,000 cells/well, and H2030 at 2,500 cells/well. 72 h after seeding, caspase-3/7 activity was measured similarly.

### Immunoblotting

For western blot analysis, parental NSCLC cells were collected at 80-90% confluency. For control non-targeting and QPRT knockdown conditions, the cells were collected either 3 days after transduction (same day for the NAD^+^ measurements) or 5 days after transduction. Cells were collected and lysed with RIPA buffer (150 mM NaCl, 1% Nonidet P-40 (IGEPAL), 0.5% sodium deoxycholate, 0.1% SDS, and 25 mM Tris) supplemented with protease inhibitors (Aprotinin 2 µg/mL; Leupeptin 10 µg/mL; Pepstatin A 1 µg/mL; PMSF 1 mM) and phosphatase inhibitors (Sodium Floride 10mM; Sodium Orthovanadate 2 mM) on a rotator at 4°C for 30 min. The supernatants of lysates were collected by centrifugation at 18,000 g x 15 min at 4°C. Protein quantification was performed using DC protein assay kit II (Biorad, 5000112). A total of 20 μg protein was used from each sample to perform SDS-PAGE under reducing condition, and the proteins were transferred onto nitrocellulose membranes (Cytiva Amersham, 10600000) via electrophoretic transfer method. For all antibodies except for anti-Caspase-3, membranes were blocked with 5% milk in TBST for 1 hour at room temperature and then incubated overnight with 5% milk in TBST using specific primary antibody. For anti-Caspase-3 rabbit polyclonal Ab (Cell Signaling Technology, 9662S) 5% BSA in TBST was used to block the membranes instead of milk. The primary antibodies used in this study were as follows: Anti-QPRT mouse monoclonal Ab (Sigma Aldrich, WH0023475M1), Anti-NAPRT rabbit polyclonal Ab (Atlas antibodies, HPA023739), Anti-PBEF/NAMPT rabbit monoclonal Ab (Cell Signaling Technology, 86634S), Anti-NADSYN1 rabbit polyclonal Ab (Atlas Antibodies, HPA038523), Anti-IDO1 rabbit polyclonal Ab (Atlas Antibodies, HPA023072), Anti-L-Kynurenine Hydrolase rabbit polyclonal Ab (Abcam, ab225916), Anti-Kynurenine 3 monooxygenase rabbit polyclonal Ab (Proteintech, 10698-1-AP), Anti-CCBL1/KAT1 Rabbit Polyclonal Ab (Proteintech, 12156-1-AP), Anti-KAT II (G-4) mouse monoclonal Ab (Santa Cruz Biotechnology, sc-377158), Anti-V5-Tag (D3H8Q) Rabbit monoclonal Ab (Cell Signaling Technology, 13202S), Anti-Caspase-3 rabbit polyclonal Ab (Cell Signaling Technology, 9662S), Anti-HAAO rabbit polyclonal Ab (Proteintech, 12791-1-AP), Anti-LC3B rabbit polyclonal antibody (Cell Signaling Technology, 2775), Anti-beta Actin Mouse monoclonal Ab (Cell Signaling Technology, 3700S). Following the primary incubation overnight, the membrane was washed 3 times for 15 min with TBST and then incubated with secondary Ab (1:10,000) for another. The blots were washed 3 times for 15 min with TBST before developing the signal. Secondary antibodies used were as follows: Anti-Mouse IgG donkey Ab, IRDye 680RD (LI-COR, 926-68072) or Anti-Rabbit IgG donkey Ab, IRDye 800CW (LI-COR, 926-32213), Anti-Mouse IgG (H+L) goat HRP Secondary Ab (Jackson ImmunoResearch, 115-035-003), Anti-Rabbit IgG (H+L) goat HRP Secondary Ab (Jackson ImmunoResearch, 111-035-003). The signal was measured using the Odyssey CLx imager (LI-COR) when IRDye secondary antibodies were used, or with Chemidoc Imaging System (Biorad) when HRP antibodies were used.

### Lysate preparation from mouse lung and tumor tissues

Established NSCLC tumors were dissected from lungs harvested from Kras^LSL-G12D/+^; Trp53^flox/flox^ (KP) (27) and Kras^LSL-G12D/+^; Lkb1^flox/flox^ (KL) mice (28), and immediately frozen in liquid nitrogen. These dissected lung tumors along with normal lung tissue from non–tumor-bearing littermate controls were prepared for western blot analyses. Frozen tissues were first pulverized using a mortar and pestle under liquid nitrogen, then further homogenized in RIPA buffer supplemented with protease and phosphatase inhibitors with ceramic beads using a bead-based homogenizer. The lysates were incubated on a rocker at 4°C for 1 hour, and total protein concentrations were measured using DC protein assay kit II (Biorad, 5000112) and lysates were analyzed as described in the immunoblotting section.

### QPRT overexpression in NSCLC cells

pLV-based control or QPRT-V5 overexpression constructs were obtained from Vector Builder (and K139A mutant version was generated in-house as described below). pMD2.G (Addgene, 12259) and psPAX2 (Addgene, 12260), PMDLg/pRRE constructs were co-transfected with each construct into HEK293T cells using the X-tremeGENE (Roche) reagent according to the manufacturer’s protocol. Media containing the lentivirus was collected as described in the previous section. To generate QPRT N-terminal overexpressed cells, the cells were transduced with the lentivirus using the spinfection method as follow: viral supernatant was added to the cell, and the cells were centrifuged for 2 hours at 750 x g. 24 h after transduction, PC9 cells were selected with 2.5 µg/mL of blasticidin (RPI, 3513-03-9), H2009 and H2030 cells with 5 µg/mL. Cells were maintained with blasticidin for the duration of the experiments to prevent drift. The expression was confirmed with Western blot analysis.

### Co-immunoprecipitation (Co-IP) with V5 tag

PC9, H2009 and H2030 cells overexpressing QPRT with an N-terminal V5 tag (pLV [Exp]-Bsd-EF1A>V5/hQPRT, Vector Builder) were cultured in 2 x 15cm^2^ dishes at 37 °C with 5% RPMI 1640 until the cells became 80-90% confluent. The medium was removed and cells were washed twice with cold PBS. 300 µL of pre-chilled low-stringency IP lysis buffer (0.05% NP-40, 50 mM NaCl, 0.5 mM EDTA, 50 mM Tris-HCl, pH 7.4) was added on each plate. The buffer was supplemented with protease/phosphatase inhibitors as described above. The cells were scraped, and the lysates were collected. The lysates from two plates were pooled together and then incubated in a rotating shaker at 4°C for 1 hour. After incubation, the tubes were centrifuged at 18,000 g, 4°C for 15 min. The supernatant containing proteins was collected, and protein concentration was determined with the DC protein assay kit II (BioRad. For inputs, 50 µg total protein were mixed with 10 µL 5x SDS-PAGE buffer and diluted to a total of 50 µL with IP lysis buffer. 20 µg protein from the input was loaded as a control. For the IP, 1 mg of total protein was diluted with IP lysis buffer to 1 mL. Anti-V5 agarose resin (Sigma Aldrich, A7345) was used as follows: the 50% bead slurry was mixed thoroughly, and 50 µL per IP was transferred to a fresh tube, followed by centrifugation at 10,000 × *g* for 30 seconds at 4°C. Then the beads were washed four times for 1 min at 1,000 x *g* with 1 mL lysis buffer. Following the final wash, resin was resuspended in an equal volume of lysis buffer to regenerate a 50% slurry. Anti-V5 agarose beads (50 µL of 1:1 slurry) were added to each lysate (1 mL) and incubated for 3 hours at 4°C on a rotator, after which the tubes were centrifuged for 5 min at 1,000 x *g* for 5 min at 4 °C. The supernatant was discarded, and the beads were washed four times for 25 min with 1 mL of IP lysis buffer at 4°C. Following the final wash, the supernatant was aspirated, and the protein-bound complexes were incubated with 50 µL of 250 µL/mL V5-peptide in PBS (Sigma-Aldrich, V7754) for 30 min on an Eppendorf thermomixer (800-900 RPM) to elute QPRT. The samples were centrifuged at 1,000 x g for 5 min at 4°C and the supernatant was collected. A second elution was performed in the same manner, and the eluates from both repeats were pooled together to a final volume of 100 µL. Eluates were analyzed by immunoblotting as described above.

### Immunohistochemistry (IHC)

Paraffin-embedded tissue sections were deparaffinized in xylene (2 × 5 min), and the slides were rehydrated through an ethanol series as follows: 2 x 100% for 3 min, 2 x 95% for 3 min, and 2 x 70% for 3 min, followed by a 2 min rinse in tap water. Antigen retrieval was performed in 10 mM citrate buffer, pH 6.0 using a pressure cooker for 15 min. After antigen retrieval, the slides were cooled down for 20 min. Then, tap water was added to the container for 5 min while it overflowed. To inhibit endogenous tissue peroxidases and reduce non-specific background, slides were incubated in 3% hydrogen peroxide (Fisher Scientific, BP2633500) diluted in tap water for 5 min. Following this step, the slides were incubated with TBST. The slides were placed on a cover plate (Epredia, 72110017) and then loaded into the Thermo-scientific Shandon Sequenza Slide Racks (Epredia 73310017) with the label up. After loading the slides, 500 µL TBST was added twice until it flowed through the rack. The slides were incubated for 20 min with 2.5% blocking reagent (goat serum, 125 µL). Primary anti-QPRT rabbit antibody (Sigma, SAB1410425) in 2.5% goat serum (1:200) in TBST was added to the slides overnight at 4°C. The next day, the slides were washed twice with TBST for 2 min, and then then incubated for 30 min with ImmPRESS HRP Goat Anti-Rabbit IgG Polymer Detection Kit, Peroxidase reagent (Vector Laboratories, MP-7451-15). After that, the slides were washed twice for 5 min with TBST. The slides were then incubated for 2 min with DAB substrate solution (1 drop of DAB solution/mL of diluent, Vector Laboratories, SK-4105). The slides were rinsed in tap water once before counterstained with hematoxylin. Hematoxylin (Vector Laboratories, H-3404-100) was added for 10 sec and slides were immediately transferred to a container with tap water to wash the excess hematoxylin. The slides were dehydrated in the alcohol series as follows: 2 x 95% for 3 min. 2 x 100% for 3 min and 2 x xylene steps for 5 min. Finally, the slides were mounted inside a fume hood using mounting medium (Vector Laboratories, H-5700-60).

### Grading of lung adenocarcinoma with simultaneous segmentation by an artificial intelligence (GLASS-AI)

Whole slide images (WSIs) were generated from H&E and immunostained slides using an Aperio ScanScope AT2 Slide Scanner (Leica) at 20x magnification with a resolution of 0.5022 microns/pixel. WSIs of immunostained sections were aligned to adjacent H&E-stained sections by a combination of global and local registration in MATLAB, first with rigid and then affine transformations, as previously described (29). The H&E-stained WSIs were then analyzed using GLASS-AI (v2.0.0; <u>github.com/jlockhar/GLASS-AI</u>), a machine learning tool for pixel-level tumor grading and segmentation in mouse models of lung adenocarcinoma(29). Qprt IHC staining was analyzed using TissueStudio (Definiens) to quantify the staining intensity within the cytoplasm of each cell in the H&E-registered WSI. The individual cells were then classified based on the GLASS-AI output of the adjacent H&E image using MATLAB (v1.0.0; <u>github.com/jlockhar/Stained-GLASS-AI</u>). The distribution of Qprt stains across different cell classes was analyzed using R Statistical Software (v4.4.2; R Core Team 2024).

### NSCLC patient analysis

QPRT gene expression data from NSCLC patients were obtained from the Lung Cancer Explorer (LCE) web portal (30), which aggregates and z-score normalizes expression data from multiple independent publicly available datasets. Data from 12 datasets comprising a total of 1,166 patients were stratified by tumor grade (Grade 1, 2, and 3). Since expression values were derived from independent cohorts profiled on different platforms, inter-dataset batch effects were corrected using the removeBatchEffect function from the limma package (31) in R. Tumor grade was included as a covariate in the design matrix to preserve biological variation of interest during batch correction. Statistical differences in QPRT expression across tumor grades were assessed using the Kruskal-Wallis test, followed by Dunn’s post-hoc pairwise comparisons in Prism.

### Orthotopic NSCLC models and bioluminescence imaging

To generate the GFP-Luciferase-labeled H2030 or PC9 cells were transduced with Luc-P2A-GFP-hygro construct (32). The cells were selected with hygromycin (100 µg/mL) until the cells in non-transduced control plate were killed. Cells were sorted based on GFP expression and subsequently maintained in RPMI 1640 media supplemented with 5% FBS. The QPRT knockdown of cells was performed as previously described. Two days after transduction, 2 × 10⁵ GFP-Luciferase-labeled H2030 or PC9 cells expressing either shQPRT or shNT were injected via the tail vein into NSG mice. One-hour post-injection, lung seeding efficiency was assessed by measuring bioluminescence by intraperitoneal injection of 150 µL luciferin (Goldbio, LUCK-2G). The IVIS 200 imaging system was used to detect the signal and take images 6 min later. Endpoint *ex vivo* imaging of lungs was performed using the same system to assess tumor burden. For endpoint *ex vivo*, the lungs were bathed in 150 µL luciferin solution for 1 min before developing the images with IVIS 200 system. The *ex vivo* imaging was performed 11 weeks after injection for H2030 and 4 weeks for PC9. For each bioluminescence image with multiple lungs, signal from the bottom right corner was taken as background. If the signal was positive, it was subtracted from the total flux signal for each lung. If the background signal was negative, then no subtraction was applied. If a final total Flux value was negative, it was reported as zero.

### QPRT site-directed mutagenesis for cell-based studies

The K139A mutant of the QPRT with a N-terminal V5 tag (pLV [Exp]-Bsd-EF1A>V5/hQPRT, Vector Builder) was generated using the QuikChange II Site-Directed Mutagenesis Kit (Agilent Technologies, 200521) according to the manufacturer’s protocol. Briefly, PCR reactions were set up using 15 ng of dsDNA plasmid template, 125 ng of each mutagenic primer, 1 μL of dNTP mix, and 1 μL of PfuUltra HF DNA polymerase (2.5 U/μL) in a 50 μL reaction volume with 1x reaction buffer. Thermal cycling was performed as follows: initial denaturation at 95°C for 30 s; 16 cycles of 95°C for 30 s, 55°C for 1 min, and 68°C for 10 min (1 min per kb of plasmid length). Following amplification, reactions were cooled on ice for 2 minutes, and 1 μL of DpnI (10 U/μL) was added to digest the parental methylated plasmid DNA. Samples were incubated at 37°C for at least 1 hour, then transformed into super competent E. coli cells for propagation. DNA from selected colonies was isolated (Omega) and the construct DNA was sent to Eton Bioscience for sequence confirmation.

### Tryptophan deprivation and viability assay

To assess the impact of tryptophan availability on QPRT-dependent cell viability, NSCLC cell lines were cultured in tryptophan-depleted medium (Caisson Labs, RPL22-6X500ML). QPRT knockdown cells were generated as described earlier. One day after transduction, H2030 cells were selected with puromycin and seeded at 1,500 cells/well in 96-well plates in normal FBS. The day after, the medium was changed to RPMI 1640 medium lacking L-tryptophan and supplemented with dialyzed fetal bovine serum (Sigma-Aldrich F0392), to remove residual amino acids. For restricted conditions, L-tryptophan was added back at a final concentration of 0 mg/L or 1.25 mg/L, compared to the normal level of 5 mg/L in RPMI 1640. Cells were maintained in these media for 72 hours before the cell death assay was performed.

### QPRT cloning, mutagenesis, and bacterial purification

The synthetic cDNA of *Homo sapiens* QPRT (*hsQPRT)* was purchased as a gBlock Gene Fragment (IDT, Coralville, IA, USA) and cloned into a pET28a plasmid via isothermal assembly. To generate hsQPRT mutants, synthetic primers were designed using the in-Fusion Cloning Primer Design Tool (Takara). All QPRT mutant constructs were verified by direct Sanger sequencing (Genewiz/Azenta Life Sciences) using T7 and T7 reverse primers. Sequence chromatograms were analyzed using SnapGene (Dotmatics) to confirm the presence of intended mutations and the absence of off-target modifications. Verified plasmids were used for subsequent expression and functional characterization.

For large-scale protein expression, plasmids containing wild-type or mutant hsQPRT cDNAs were transformed into BL21 DE3 cells (NEB). Transformed cells were grown in LB medium supplemented with 50 µg/L kanamycin until the optical density (OD_600_) reached 0.6–0.8. Protein expression was induced by adding 500 µM IPTG, followed by incubation at 18°C for 20 hours. Induced cells were harvested by centrifugation and resuspended in harvest buffer (20 mM Tris, pH 7.5, 500 mM NaCl, 25 mM imidazole, 5 mM BME). For protein purification, the resuspended cells were lysed by sonication and centrifuged at 40,000 × g for 30 minutes at 4°C to isolate the soluble fraction. The supernatant was loaded onto a 10 mL Ni²⁺ affinity column pre-equilibrated with Nickel A buffer (20 mM Tris, pH 7.5, 300 mM NaCl, 25 mM imidazole, 5 mM BME). The protein was eluted using a gradient of Nickel B buffer (20 mM Tris, pH 7.5, 300 mM NaCl, 500 mM imidazole, 5 mM BME). The eluted protein was concentrated to 1 mL using a 30 kDa molecular weight cutoff (MWCO) filter (Millipore Sigma) and further purified by size-exclusion chromatography (SEC) on a HiLoad Superdex 200 16/60 column, equilibrated with SEC buffer (20 mM HEPES, pH 8.0, 150 mM NaCl, 2 mM DTT). Fractions containing purified hsQPRT were concentrated to 4 mg/mL, and flash-frozen in liquid nitrogen for long-term storage at −80°C. Fractions were analyzed by SDS-PAGE on 4 – 15% Mini-PROTEAN TGX Stain-Free gels (Bio-Rad, #4568086). Protein bands were visualized using Stain-Free detection on a ChemiDoc MP system (Bio-Rad).

### QPRT *in vitro* enzymatic activity

The continuous kinetic assay of QPRT activity to screen QPRT mutants was performed using the PiPer™ Pyrophosphate Assay Kit (Thermo Fisher Scientific, P22062) in a 96-well microplate format. The PiPer™ detection reagent, containing Amplex® Red, horseradish peroxidase (HRP), maltose phosphorylase, glucose oxidase, and inorganic pyrophosphatase, was prepared according to the manufacturer’s instructions. The enzyme reaction was initiated by mixing equal volumes of the PiPer™ detection reagent and QPRT at a final concentration of 5 µM, in the presence of 20 µM QA and 100 µM phosphoribosyl pyrophosphate (PRPP) in PiPer™ reaction buffer. The pyrophosphate-coupled formation of resorufin was continuously monitored by measuring absorbance at ∼560 nm every 2 minutes using a multi-mode plate reader, Agilent Biotek Synergy H1. Appropriate controls, including reactions lacking either enzyme or substrate, were included to correct for background signals.

### QPRT sequence conservation analysis and molecular visualization

Amino acid conservation analysis of hsQPRT was performed using ConSurf (33) to assess evolutionary conservation at individual residue positions. The structural model used for mapping conservation scores was retrieved from the Protein Data Bank (PDB ID: 5AYY). The ConSurf server was run with default parameters, utilizing a homologous sequence dataset obtained from UniProt and multiple sequence alignment via Clustal Omega. Conservation scores were color-coded from variable (cyan) to highly conserved (maroon) and mapped onto the QPRT structure. For structural visualization and figure generation, PyMOL (Schrödinger, LLC) was used to render high-resolution molecular images. The conservation scores from ConSurf were loaded into PyMOL as a B-factor color gradient.

### Stable isotope tracer analysis

PC9, H2009, H2030, H1650, H2122, H1944, KP 1.9 and KL 95 NSCLC cell lines were treated with d_4_-NAM (32 µM, Cambridge Isotope Laboratories, DLM-6883-0.1), ^13^C_6_-NA (20 µM, Cambridge Isotope Laboratories, CLM-9954-0.001), d_3_-QA (20 µM, CDN Isotopes, D-8099) and ^13^C_11_-Trp (80 µM, Cambridge Isotope Laboratories. CLM-4290-H-0.1) for 24 h. The cells were washed with ice-cold PBS twice, scraped using a cold cell-lifter, and centrifuged at 300 x g for 5 min. The supernatants were discarded, and the pellets were snap frozen and stored at −80°C for further analysis. KP mice were either left uninfused (control), or continuously infused via jugular catheter for 4 hours with one of the following stable isotope tracers, each prepared in sterile saline: d_4_-NAM (4 mM), ^13^C_6_-NA (0.2 mM), d_3_-QA (4 mM) or ^13^C_11_-Trp (100 mM). All infusions were conducted at a rate of 120 µL/hr, with the uninfused group serving as a shared control across all tracer conditions. Following infusion, tumors were dissected and snap frozen. Graphs in Fig. 3C, E, G, I, and Supplementary Fig. S2C, E, G, I all utilized the data from this single “no infusion” group as the control.

Frozen tissue samples were processed for NADome metabolites as described before(34,35). Briefly, tissues were weighed and homogenized in ice-cold 1:1 methanol (MeOH):water (400 mL per 20 mg tissue weight) using a liquid nitrogen-chilled tissue homogenizer (Precellys). NADome metabolites from NSCLC cell lines and tissue lysates were extracted with 400 µL of 80% (1:1) MeOH:acetonitrile (ACN) + 0.1% formic acid followed by neutralization with 15% (w/v) ammonium bicarbonate and centrifugation at 18,000 x g for 30 min at 4°C. The supernatants were dried under nitrogen and resuspended in 50 µL of 20 mM ammonium bicarbonate (pH 8.0) for analysis using a QExactive Plus Orbitrap MS (QE-MS) with a HESI II probe and coupled with an Ultra High-Pressure Liquid Chromatography (Dionex UltiMate 3000) system. Samples were analyzed within 24 h of reconstitution. Chromatographic separation of NADome metabolites was performed using reverse-phase (RP) chromatography HSS C18 column (150 x 2.1 mm i.d., 1.7 mm; Waters) with a flow rate of 0.4 mL/min. Solvent A was 0.1% formic acid in water and solvent B was 0.1% ACN. The LC gradient included a 1 min hold at 0.1% solvent B followed by a ramp from 0.1% to 50% solvent B over the next 7 min followed by a ramp to 99% over the next 8 min. A 3 min hold at 99% was followed by a return to 0% solvent B over the next 0.5 min. The run was completed with a 7 min recondition at 0.1% solvent B. The RP separation was performed over 22 min in positive ionization mode. Nucleotide separation was achieved with BEH amide column (150 x 2.1 mm i.d., 1.7 mm; Waters). The column temperature was set at 30°C, and the injection volume was 5 mL. Solvent A is 20 mM ammonium bicarbonate pH 9.5 and solvent B is ACN. The mass spectrometer was operated in positive and negative ESI ion mode in the scan range of m/z 70–1,050 with the resolution of 70,000 at m/z 200, automatic gain control (AGC) target at 1 3 106 and maximum injection time of 50 ms. The spray voltage was set to 3.5 and 2.5 kV in positive and negative ion mode, respectively. The heated capillary was set at 200°C; the HESI probe was set at 350°C; and the S-lens RF level was set at 45. The gas settings for sheath, auxiliary and sweep were 40, 10 and 1 unit, respectively. Peak areas (AUC) were used for comparative quantitation. RAW data files were processed with Xcalibur Quan software (Thermo Scientific). The identity of each mass metabolite was confirmed with the respective authentic standard, and the data were corrected for natural isotopic abundance. An in-house library containing all possible 2H and 13C isotopologues (m/z of M0, M1, M2, M3.Mn) of each relevant metabolite was used for the assignment of AUC of LC-MS signals. Fractional enrichment for each targeted metabolite is an index of the relative flux as calculated from the MID profile using a linear simultaneous equation.

### Statistical Analysis

Statistical analyses were performed either with GraphPad Prism or with R, and a p value of <0.05 was considered statistically significant. For each figure, sample sizes and statistical tests are listed in the figure legends. Unless otherwise indicated, an unpaired Student’s *t*-test was performed between 2 groups and ANOVA between multiple groups, and graphical data were represented as mean +/- SEM.

### Data Availability

All data can be made available upon reasonable request to the corresponding author at.

## Results

### QPRT levels are elevated in NSCLC and correlate with NSCLC progression

Several studies put forward NAD^+^ levels as major regulators of the metabolic and redox reprogramming that empowers NSCLC progression (6,36–38). Considering that bypassing the reliance on the NAD^+^ salvage pathway is a characteristic of most cancer cells (6), we hypothesized that alternative NAD^+^ biosynthetic pathways might play an important role in NSCLC progression. To gain insights into this possibility we first evaluated the levels of the rate-limiting enzymes of the different NAD^+^ biosynthetic pathways (Supplementary Fig. S1A), including NAMPT (salvage), NAPRT (Preiss-Handler) and QPRT (*de novo*), in a panel of NSCLC cell lines that recapitulates the diversity of oncogenic drivers of NSCLC, including mutant KRAS and EGFR. Strikingly, even though all 3 enzymes were expressed in these cells to varying degrees, QPRT levels were more consistently elevated than the levels of NAPRT and NAMPT in this NSCLC cell line panel (Fig. 1A). To assess the physiological relevance of these findings *in vivo* we dissected lung tumors from well-established genetically engineered mouse models of NSCLC, the KRAS/p53-driven (KP) (39) and KRAS/LKB1-driven (KL) (28) models, and compared the expression of these enzymes in the NSCLC tumors to those in non-tumor bearing normal lung tissue. In line with what was observed in human NSCLC cell lines, QPRT was highly expressed in both KP and KL tumors. Importantly, while NAMPT and to some extent NAPRT were also expressed in the normal lung, albeit in lower levels, QPRT’s expression in the normal lung was minimal (Fig. 1B). This suggested that QPRT expression is a feature that arises in the NSCLC progression continuum. To directly assess this possibility, we evaluated QPRT levels in the lungs of the KP and KL models by immunohistochemistry (IHC). Further analysis revealed a correlation between QPRT levels and the grade of the NSCLC tumors across both models, where higher grade tumors exhibited higher QPRT expression (Fig. 1C&D), a feature that is retained in patient samples (Fig. 1E). Together, these data point to a critical role for QPRT in NSCLC progression, whose levels rise upon NSCLC establishment and continue to increase as these tumors progress into higher grades.

**Figure 1.**
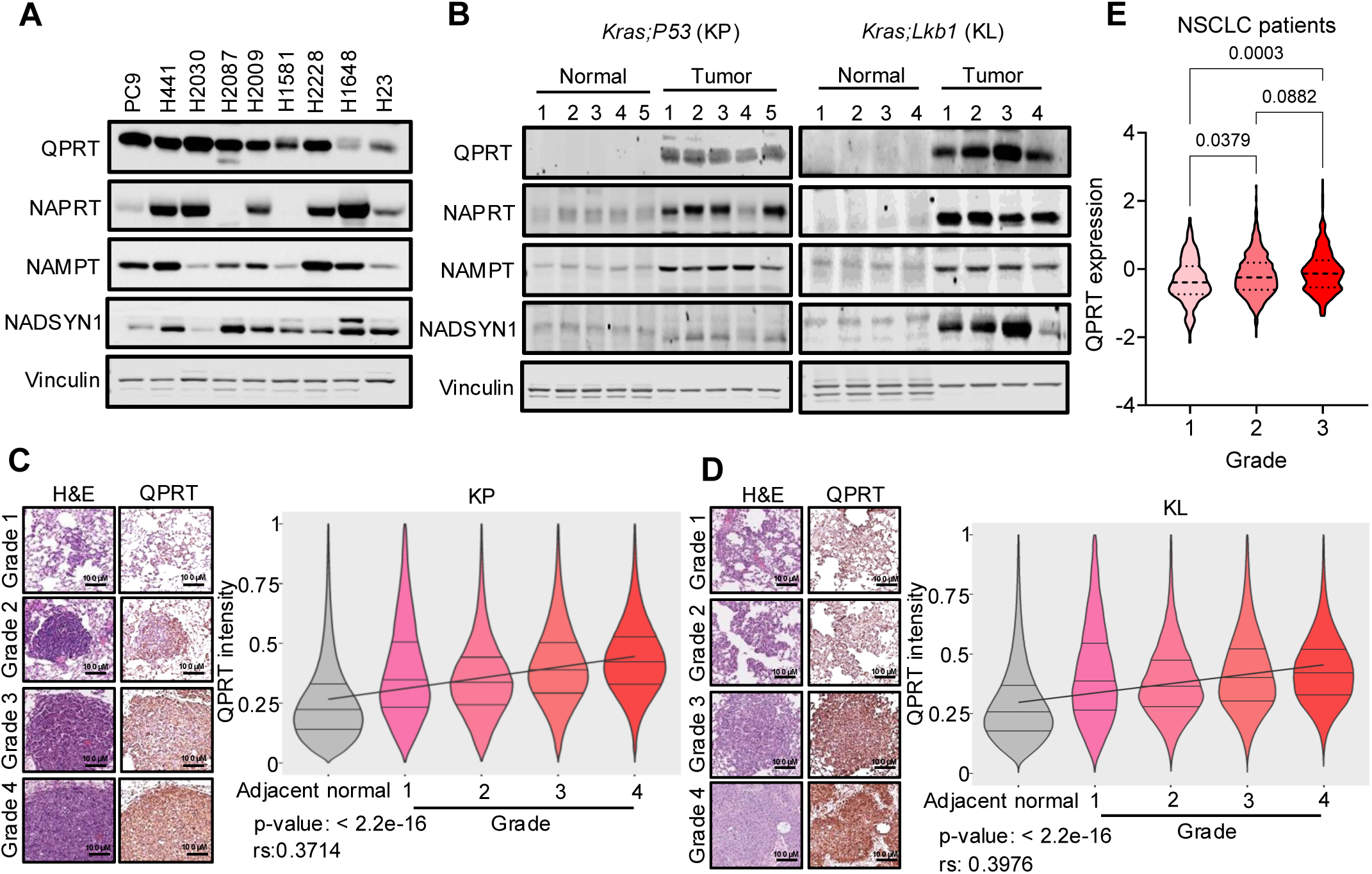
QPRT is upregulated in NSCLC cell lines and NSCLC tumors in genetically engineered mouse models. **(A, B)** Immunoblot analysis of QPRT and other NAD⁺ biosynthetic enzymes in a panel of NSCLC cell lines (representative images, n = 4) **(A),** and NSCLC tumors from KP and KL GEM models (KP: Kras^LSL-G12D/+^; Trp53^flox/flox^, KL: Kras^LSL-G12D/+^; Lkb1^flox/flox^: n=5 for KP, n = 4 for KL and n=5 normal lung tissue from non–tumor-bearing mice) **(B)**. **(C, D)** Representative IHC images and quantification of QPRT protein levels in IHC slides across different tumor grades using GLASS-AI in KP **(C)** and KL **(D)** GEMM models (n = 5 mice for each). For D and E, the statistical analysis was performed using Spearman correlation. **(E)** Violin plots showing z-score of QPRT expression in Grade 1, Grade 2, and Grade 3 NSCLC tumors (n = 1,166 patients) after batch correction. Statistical comparisons between grades were performed using the Kruskal-Wallis test followed by Dunn’s post-hoc pairwise comparisons.

### Loss of QPRT suppresses tumor growth in NSCLC by inducing cell death

Having demonstrated that high QPRT levels are a feature of NSCLC that is associated with NSCLC progression into high-grade tumors (Fig. 1C-E), we sought to determine its functional relevance in NSCLC. To test this, we knocked down QPRT in several NSCLC cell lines (PC9, H2009, H2030 and H1581), which express high levels of QPRT (Fig. 1B) and evaluated its consequences for their growth. Remarkably, QPRT knockdown (Fig. 2A) abolished the ability of these cells to grow *in vitro* in 2D (Fig. 2B) and in 3D as spheroids (Fig. 2C&D), suggesting that QPRT suppression might hinder the growth and progression of NSCLC tumors. To directly test this premise, we transplanted luciferase-expressing QPRT-suppressed PC9 and H2030 cells into the lungs of immunocompromised mice. In line with the *in vitro* data, the suppression of QPRT significantly abrogated tumor growth in both models (Fig. 2E&F) despite having no effect in the ability of these cells to seed in the lungs (Supplementary Fig. S1B). To elucidate the QPRT-driven processes that underlie its pro-tumorigenic effects in NSCLC, we first evaluated if QPRT suppression affects proliferative capacity by tracking the levels of the fluorescent dye DiD, which uniformly labels cell membranes and whose abundance decreases upon cell division (Supplementary Fig. S1C) (40). DiD retention was higher in some conditions but was inconsistent across shRNAs and cell lines (Supplementary Fig. S1D). To further investigate the effects of QPRT on NSCLC proliferation, we evaluated cell cycle progression in the same conditions. Similarly to what was observed with DiD retention, the effects on each cell cycle phase were inconsistent across shRNAs and cell lines (Supplementary Table 1), and therefore, do not explain the consistent inhibition of cell growth observed across cell lines upon QPRT knockdown (Fig. 2B). Tumor growth reflects the net balance between cellular proliferation and cell death (41). Considering the inconsistent effects of QPRT suppression on NSCLC proliferation, we reasoned that QPRT loss might suppress tumor growth by causing cell death. To test this, we monitored cell death upon QPRT knockdown in NSCLC cells and observed a consistent increase in cell death upon QPRT loss (Fig. 2G-I), matching the effects on cell growth (Fig. 2B). Together, these data showed that NSCLC cells rely on QPRT for survival.

**Figure 2.**
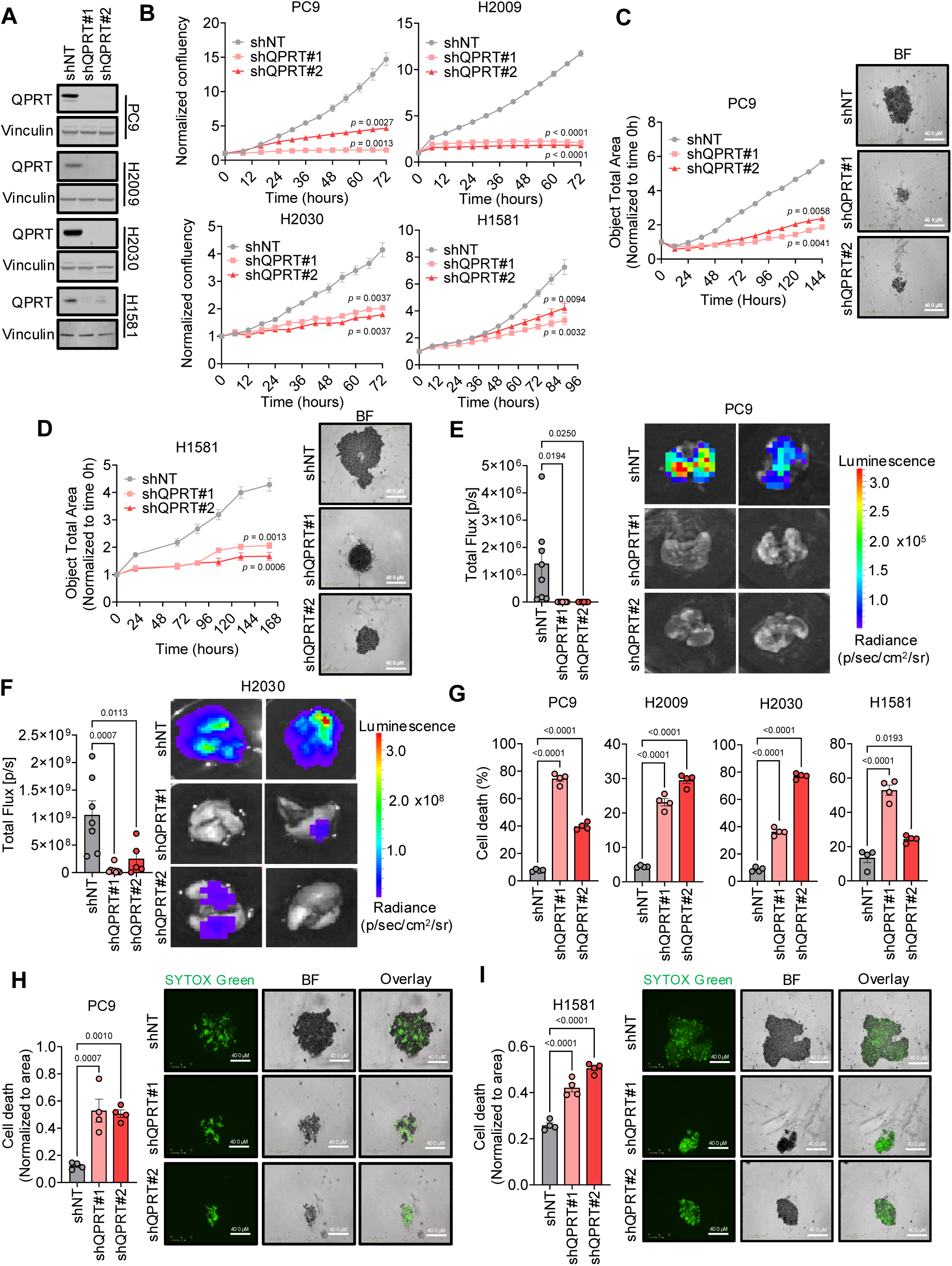
QPRT knockdown inhibits NSCLC cell growth and induces cell death *in vitro* and *in vivo.* **(A, B)** Immunoblot confirmation of shRNA-mediated QPRT knockdown in PC9, H2009, H2030, and H1581 cells (representative images) **(A)**, and quantification of 2D growth assays of these cells using Incucyte live-cell imaging analysis **(B)** (n = 4). **(C, D)** Quantification of 3D spheroid growth assays using Incucyte live-cell imaging analysis and representative end-point images of PC9 **(C)** and H1581 **(D)** cells upon QPRT knockdown (n = 4). **(E, F)** Quantification and representative images of tumor burden with *ex vivo* bioluminescence of lungs harvested from NSG mice implanted with shNT or shQPRT-expressing PC9 **(E)** or H2030 **(F)** cells (for PC9: shNT n = 8, shQPRT#1 n = 7, shQPRT#2 n = 6; for H2030: shNT n = 7, shQPRT#1 n = 8, shQPRT#2 n = 5, p/s indicates photons/second, “p/sec/cm²/sr” stands for photons per second per square centimeter per steradian). **(G)** Quantification of cell death using SYTOX Green staining in 2D cultures of NSCLC cells upon QPRT silencing (n = 4). **(H, I)** Quantification of cell death using SYTOX Green in 3D spheroid cultures and representative images for PC9 **(H)**, or H1581 **(I)** cells upon QPRT silencing (n = 4, BF: brightfield). Data are represented as the mean ± SEM (B-I). For the graphs in panels B-D statistical significance determined by two-way repeated measures ANOVA followed by Dunnett’s post hoc test and p value for last time point is included on the graphs. For E-I one-way ANOVA was performed followed by Dunnett’s post hoc test. The scale bars on representative images in C, D, H and I represent 400 µm.

### QPRT regulates cell death via a mechanism unrelated to NAD⁺ metabolism

QPRT is the rate-limiting enzyme in the *de novo* NAD^+^ biosynthetic pathway (24) and NAD^+^ levels are well-known to regulate cell fate decisions, including cell death in NSCLC (23,38,42). Consequently, we hypothesized that QPRT suppression triggers cell death by depleting NAD^+^. To test if this is indeed the case, we measured NAD^+^ levels in NSCLC cells upon QPRT knockdown and, surprisingly, observed no changes (Fig. 3A). NAD⁺ biosynthetic pathways are known to compensate for one another to maintain cellular homeostasis and support cell viability (43,44). Considering that loss of QPRT in NSCLC cells does not cause changes in NAD^+^ levels, we evaluated the levels of the remaining rate-limiting enzymes of the other NAD^+^ biosynthetic pathways. Importantly, our data showed that no elevation of either NAPRT or NAMPT occurs upon QPRT knockdown in NSCLC cells (Supplementary Fig. S2A), suggesting that no compensation is at play. To precisely evaluate the contribution of each NAD^+^ biosynthetic pathway, we traced the incorporation of labeled precursors of each of these pathways in different human and murine NSCLC cell lines *in vitro*, as well as *in vivo* using the KP model. As expected, [d_4_]-NAM, which enables evaluation of salvage pathway activity, showed label incorporation into NAD^+^ across the different cell lines as well as *in vivo* (Fig. 3B&C, Supplementary Fig. S2B&C). Likewise, the incorporation of label from [^13^C_6_]-NA into NAD^+^ was also observed in several cell lines despite not being observed in the KP model *in vivo* under these experimental settings (Fig. 3D&E, Supplementary Fig. S2D&E). This supported the idea that the main pathway of NAD^+^ generation in NSCLC cells is the salvage pathway with a smaller contribution from the Preiss-Handler pathway. However, no incorporation of label from either [d_3_]-QA, QPRT’s direct substrate (Fig. 3F&G and Supplementary Fig. S2F&G) or its precursor, [^13^C_11_]-Trp (Fig. 3H&I and Supplementary Fig. S2H&I) was observed in any of the NSCLC cell lines *in vitro* or in the KP tumors *in vivo*. These findings support the notion that NSCLC cells do not utilize QA or Trp for NAD⁺ biosynthesis under normal conditions. Inhibition of the salvage pathway *in vitro* is well known to significantly diminish NAD^+^ pools and consequently cause cell death in cancer cells (45,46). Thus, we reasoned that the *de novo* pathway might only play a role in conditions where the salvage pathway is insufficient to meet the demands for NAD^+^. Treatment of NSCLC cells with the well-established NAMPT inhibitor, FK866 (47), caused a pronounced decline in NAD^+^ levels (Fig. 3J). In line with our tracing data, treatment with either NMN or nicotinamide riboside (NR), NAD^+^ precursors that bypass NAMPT to fuel the salvage pathway (13), was sufficient to restore NAD^+^ levels (Supplementary Fig. S2J). On the other hand, neither treatment with QA nor Trp was able to recover the drop in NAD^+^ levels caused by NAMPT inhibition (Fig. 3J), supporting the idea that the *de novo* NAD^+^ biosynthetic pathway is not at play in NSCLC. In line with this, Trp restriction had no effect on NAD^+^ levels in NSCLC cells regardless of the presence or absence of QPRT (Supplementary Fig. S3E). Likewise, treatment of QPRT-knockdown NSCLC cells with NMN or NR, which induces NAD^+^ levels in these cells, had no effect on cell death (Supplementary Fig. S2K). Together, these data demonstrated that while QPRT regulates cell death in NSCLC, its effects are independent of its role in the *de novo* NAD^+^ biosynthetic pathway.

**Figure 3.**
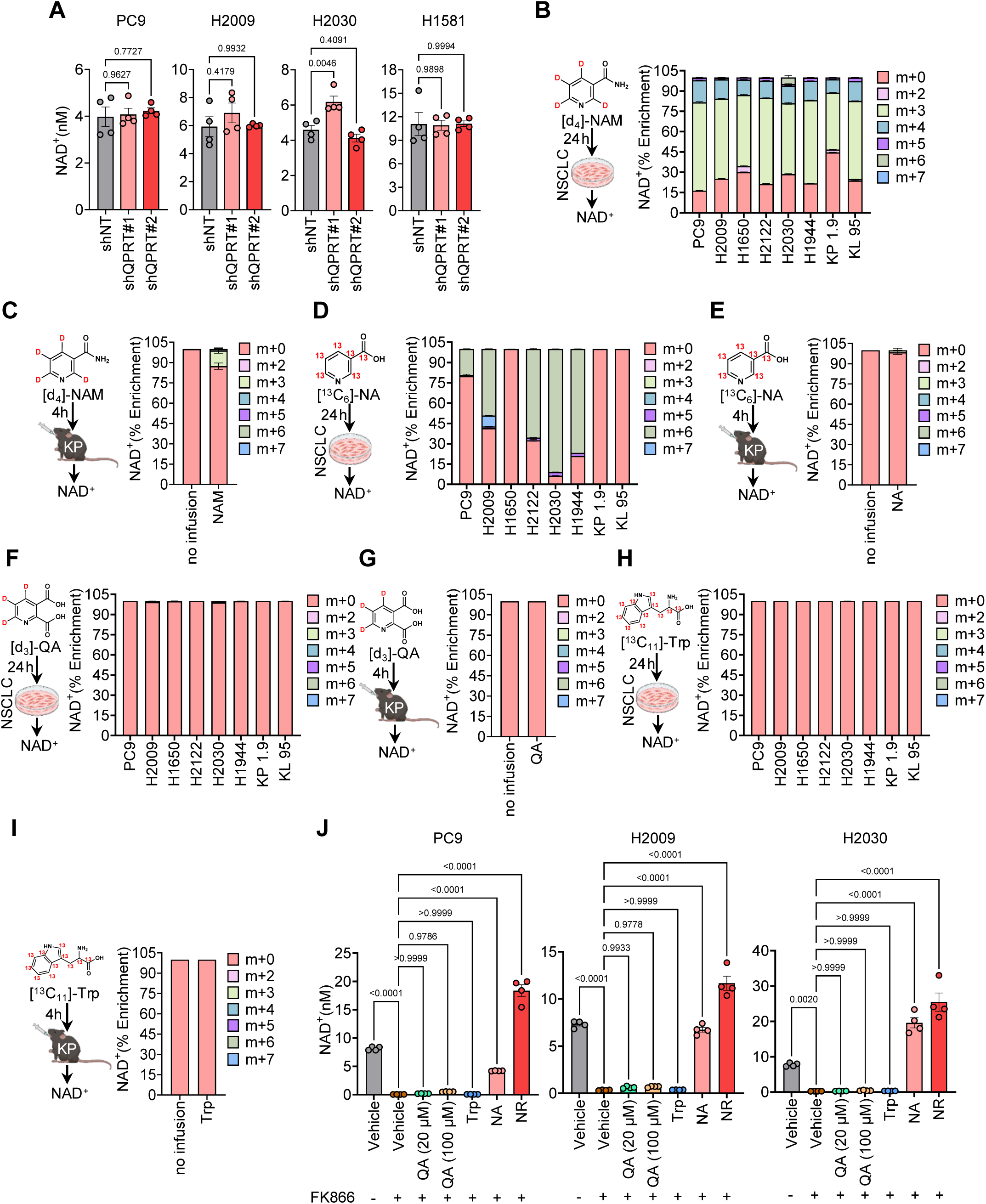
QPRT silencing-induced cell death is independent of NAD⁺ levels and *de novo* biosynthesis. **(A)** NAD⁺ quantification in NSCLC cells following QPRT knockdown (n = 4). **(B–I)** Stable isotope tracing for NAD^+^ *in vitro* in NSCLC cells after 24h treatment using d_4_NAM (32 µM) **(B)**, ^13^C_6_NA (20 µM) **(D)**, d_3_QA (20 µM) **(F)**, and ^13^C_11_Trp (80 µM) **(H)**, (n = 5) and *in vivo* in KP tumors from mice either with no infusion or after 4h infusion using d_4_NAM (4 mM) **(C)**, ^13^C_6_NA (0.2 mM) **(E)**, d_3_QA (4 mM) **(G)**, and ^13^C_11_Trp (100 mM) **(I)** (n = 5). **(J)** NAD⁺ levels in cells treated with FK866 (NAMPT inhibitor, 50 µM), with or without precursor supplementation (n = 4). Data are represented as mean ± SEM with statistical significance measured by one-way ANOVA followed by Dunnett’s post hoc test for A and Tukey’s post hoc test for J.

### QPRT modulates NSCLC survival independently of its role on tryptophan catabolism

Given our findings establishing a role for QPRT in regulating NSCLC survival independently of its role in fueling NAD^+^ and the well-characterized role of QPRT in the kynurenine pathway of tryptophan catabolism (Supplementary Fig. S3A), we hypothesized that loss of QPRT might cause cell death in NSCLC cells through metabolic disruption of this pathway and accumulation of specific metabolites upstream of QPRT. One of the main metabolic consequences of suppressing QPRT is the accumulation of QA, a well-known neurotoxin that causes cell death in neurons through hyperactivation of N-methyl-D-aspartate (NMDA) receptors (48,49). Importantly, treatment with different concentrations of QA had no cell death-promoting effects in NSCLC cells (Supplementary Fig. S3B), excluding QA accumulation as the cause of cell death upon QPRT loss. Additionally, no changes in the expression of the enzymes of the kynurenine pathway were observed upon QPRT knockdown in NSCLC cells (Supplementary Fig. S3C), suggesting that the cell death-promoting effects of loss of QPRT are also unrelated to changes in the kynurenine pathway. To determine if the effects of QPRT in regulating cell death are indeed independent of Trp catabolism, we restricted Trp availability to NSCLC cells with QPRT knockdown and evaluated the consequences for cell death. Strikingly, suppression of QPRT induced cell death in NSCLC cells independently of Trp availability (Supplementary Fig. S3D).

### QPRT regulates apoptosis in NSCLC through its interaction with caspase 3

To better understand the mechanism by which QPRT regulates cell death in NSCLC, we next investigated the mode of cell death triggered by QPRT loss. Considering that QPRT is a metabolic enzyme, we first assessed if the cell death observed upon QPRT loss is mediated by ferroptosis, an iron-dependent cell death mode triggered by the accumulation of lipid peroxides (50). Treatment of NSCLC cells with the ferroptosis inhibitor, ferrostatin 1 (51), did not prevent the induction of cell death caused by QPRT loss (Supplementary Fig. S4A) excluding ferroptosis as the mode of cell death governed by QPRT in NSCLC. Next, we evaluated whether autophagic cell death was at play. Evaluation of LC3B showed that QPRT knockdown does not induce autophagic activity in NSCLC cells (Supplementary Fig. S4B), supporting a role for QPRT in regulating a cell death mode other than autophagy. Apoptosis is a tightly regulated form of programmed cell death essential for tissue homeostasis and development (52). Importantly, in many cancers, apoptotic pathways are disrupted, allowing cancer cells to evade death signals, accumulate genetic instability and progress (53,54). Consequently, we sought to evaluate if the cell death triggered by loss of QPRT is driven by apoptosis, and evaluated the levels of Annexin V, a well-established apoptosis marker (55). Consistent with apoptosis as the mode by which QPRT regulates cell death in NSCLC, QPRT-deficient cells exhibited a marked increase in Annexin V positivity in both 2D monolayer cultures and 3D spheroid models (Fig. 4A-B). To corroborate this finding, we treated NSCLC cells with Z-VAD-FMK, a pan-caspase inhibitor known to block apoptosis (55,56). Strikingly, the presence of Z-VAD-FMK was sufficient to prevent cell death induced by QPRT in NSCLC cells (Fig. 4C) demonstrating that QPRT’s effects in NSCLC are mediated by apoptosis. In support of this premise, previous work aimed at identifying caspase 3 binding partners put forward QPRT as a regulator of spontaneous cell death by suppressing overproduction of active caspase 3 in HeLa cells (57). Thus, we reasoned that a similar mechanism might be at play in NSCLC. To interrogate this question, we first evaluated if QPRT interacts with caspase 3 in the context of NSCLC. Co-immunoprecipitation experiments revealed that QPRT interacts with caspase 3 in NSCLC cells (Fig. 4D), which correlated with increased caspase 3/7 activity in these cells upon QPRT suppression (Fig. 4E). Moreover, to ensure that this feature of QPRT is independent of its role in NAD^+^ metabolism, we treated NSCLC cells with NR to raise NAD^+^ levels and evaluated its consequences for caspase 3/7 activity. In line with QPRT regulating cell death independently of NAD^+^, treatment with NR did not consistently reduce caspase 3/7 activity induced by loss of QPRT (Supplementary Fig. S4C). Together, these data support the premise that QPRT protects NSCLC cells from spontaneous cell death by regulating caspase 3 independently of its catalytic activity and thereby empowers NSCLC progression. Structural characterization of QPRT has previously established key amino acid residues responsible for QPRT’s catalytic activity, including R138, K139, and K171 (Fig. 4F) (58,59). To discern if the regulation of caspase 3 is indeed independent of QPRT’s catalytic activity, we performed site-directed mutagenesis to generate and purify catalytically inactive mutant versions of QPRT, including QPRT^R138Q^, QPRT^K139A^, and QPRT^K171A^. Evaluation of QPRT enzymatic activity in biochemical assays using purified protein (Supplementary Fig. S4D-E) demonstrated that while QPRT^R138Q^ only minorly affects its catalytic activity, QPRT^K139A^ and QPRT^K171A^ had a striking reduction in QPRT activity with QPRT^K139A^ being catalytically dead (Fig. 4G), prompting us to focus on this mutant for subsequent studies. Strikingly, co-immunoprecipitation analyses of NSCLC cells expressing either wild-type QPRT (QPRT ^WT^) or its catalytically inactive mutant, QPRT^K139A^, showed that the interaction of QPRT with caspase 3 is retained with QPRT^K139A^ (Fig. 4H), demonstrating its independence from QPRT catalytic activity. Interestingly, this interaction is not only retained but also to some extent increased in the catalytically inactive mutant (Fig. 4H), suggesting that a switch may exist that directs QPRT towards its function in the *de novo* NAD^+^ biosynthesis pathway, which is dependent on its catalytic activity, versus its function as an apoptosis regulator independently of its enzymatic function.

**Figure 4.**
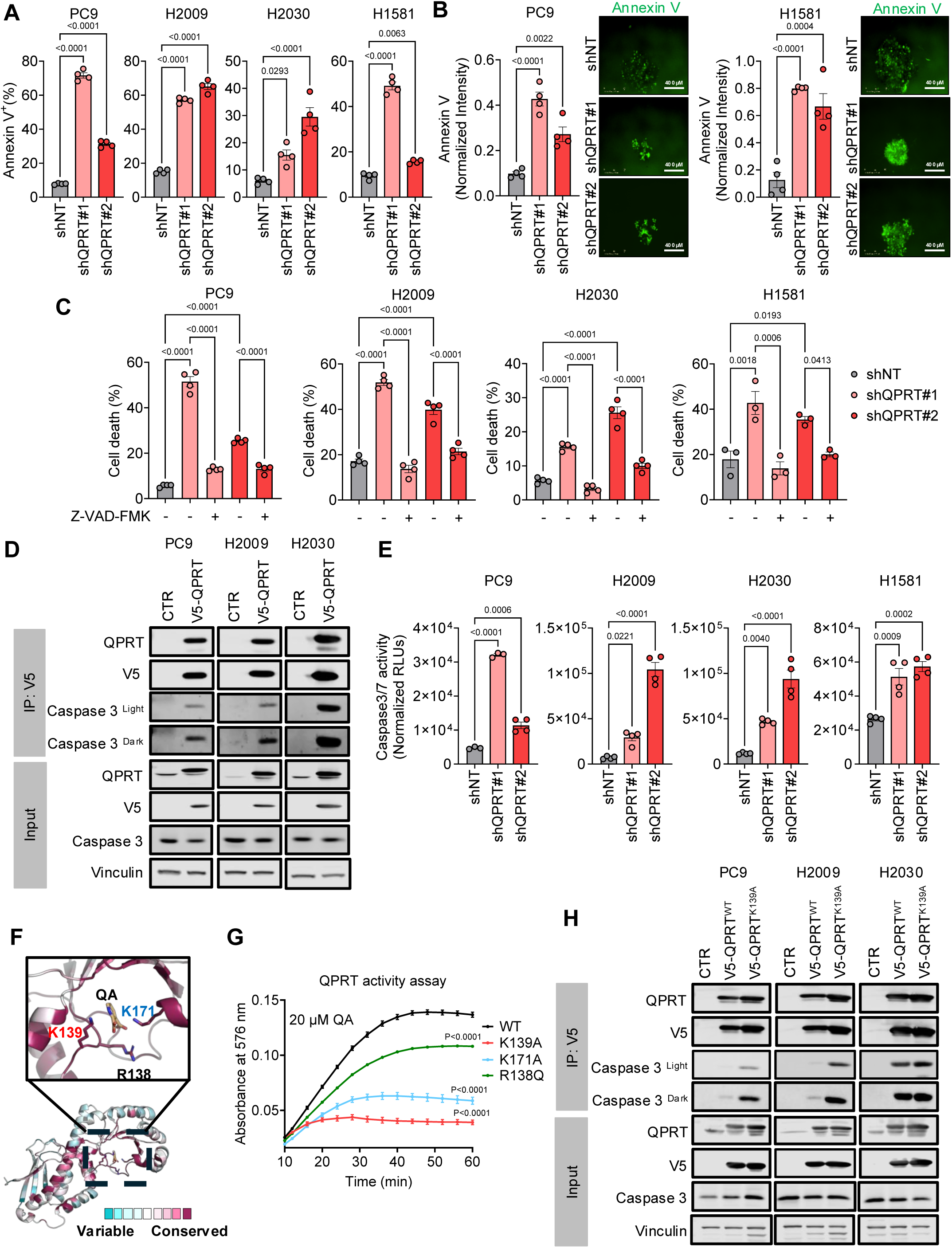
QPRT silencing induces apoptosis through non-catalytic interaction with caspase-3. **(A–B)** Annexin V, CF450 staining (0.25 µg/ml) in NSCLC cells with QPRT knockdown cultured in 2D **(A)** and 3D conditions with representative spheroid images **(B)** (n = 4). **(C)** Quantification of cell viability using PI staining for PC9, H2009 and H2030, and SYTOX Green for H1581 cells with QPRT silencing upon Z-VAD-FMK (50 µM for PC9, H2009 and H2030, and 100 µM for H1581) treatment (n = 4). **(D)** Immunoblots for co-IP experiments of V5-QPRT and its interaction with pro-caspase 3 in PC9, H2009 and H2030 cells. **(E)** Caspase-3/7 activity level in NSCLC cells with QPRT knockdown (n = 4). **(F)** Ribbon representation of QPRT (PDB: 5AYY) in complex with quinolinic acid with active site residues shown as sticks. Residues are colored according to ConSurf conservation scores**. (G)** Measurement of QPRT enzymatic activity over time of wild-type and catalytically inactive QPRT mutants purified form bacterial cells (n = 3). **(H)** Immunoblots for caspase-3 co-immunoprecipitated with catalytically inactive QPRT^K139A^ versus wild-type QPRT. Data are represented as the mean ± SEM with statistical significance measured by one-way ANOVA followed by Dunnett’s post hoc test for panels A, B, and E, and followed by Tukey’s post hoc test for panel E. For the graph in panel G statistical significance was measured by two-way repeated measures ANOVA followed by Dunnett’s post hoc test and p value for the last time point is shown on the graphs. Scale bars in B indicate 400 µm.

## Discussion

In this study, we identified QPRT as a critical regulator of apoptosis in NSCLC. This function was through a non-catalytic mechanism independent of QPRT’s canonical role in the *de novo* NAD⁺ biosynthetic pathway. Our data revealed that QPRT enzymatic function that converts quinolinic acid (QA) to nicotinic acid mononucleotide (NAMN), an NAD^+^ precursor, was dispensable for NSCLC cell survival. Metabolic tracing showed that neither QA nor its precursor Trp contributes to NAD⁺ biosynthesis in NSCLC cells under basal or stress conditions. Moreover, QPRT knockdown had no impact on NAD⁺ levels, and NAD⁺ precursors such as NMN and NR failed to rescue QPRT-depleted cells from death, ruling out a compensatory or metabolic function. Instead, we found that QPRT’s anti-apoptotic role in NSCLC is mediated by its direct interaction with caspase 3, which suppresses caspase 3 activation and thereby prevents apoptosis (Fig. 5). This observation is consistent with prior work that identified QPRT as a binding partner of caspase 3 in other cell types (57). Importantly, catalytically inactive QPRT can still interact with caspase 3, highlighting a moonlighting function that relies on a structural interaction independent of QPRT’s enzymatic activity. This new non-canonical function of QPRT challenges the current paradigm that QPRT’s relevance to cancer is primarily due to its role in NAD⁺ metabolism. It’s important to note that targeting NAD⁺ biosynthesis, particularly through NAMPT inhibition, has shown preclinical promise, but such approaches have had limited success due to toxicity and metabolic redundancy (6,16,60).

**Figure 5.**
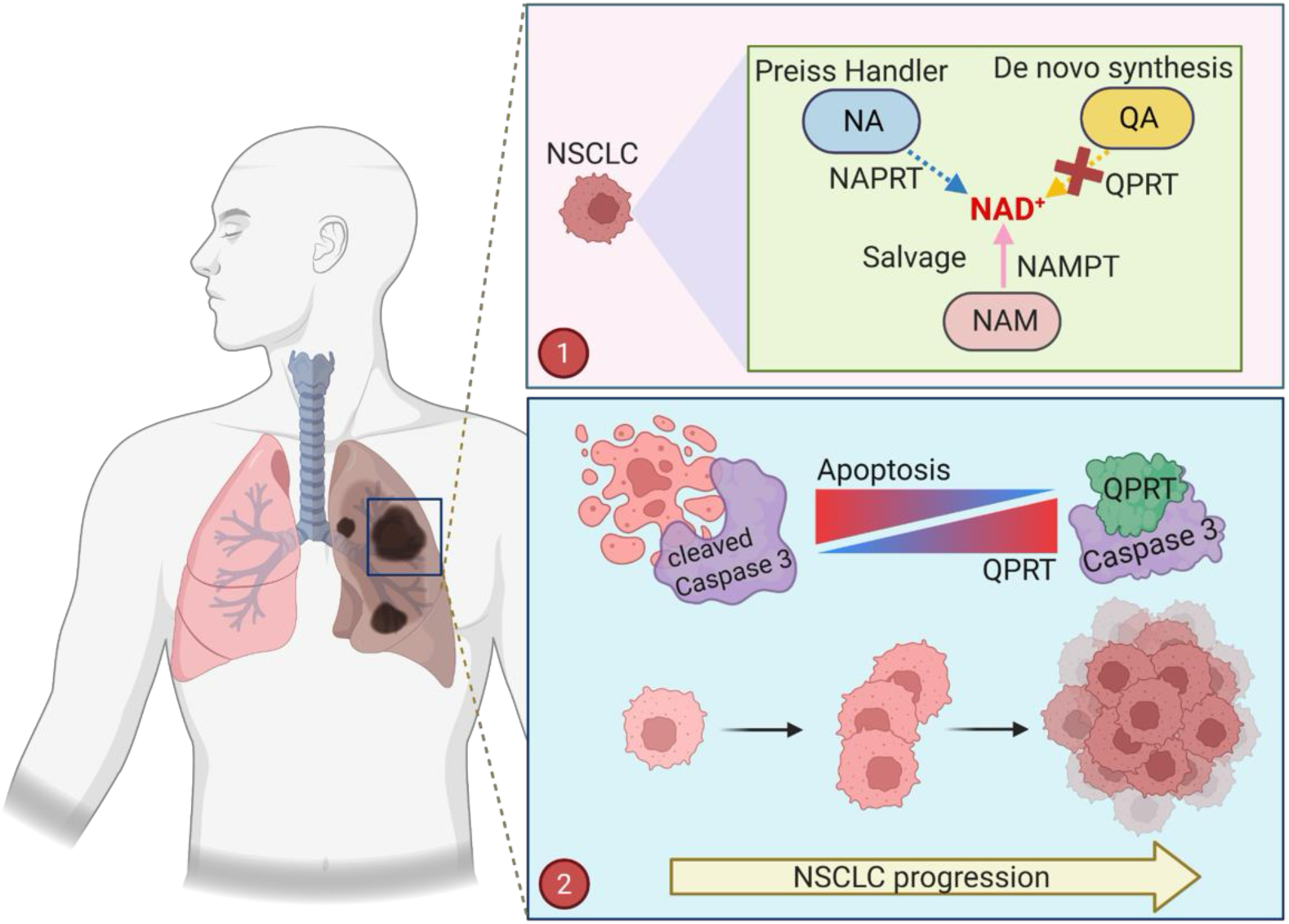
QPRT moonlights as an apoptosis regulator in NSCLC, empowering NSCLC progression independently of its enzymatic function. In NSCLC cells as diagrammed in (**1**) *de novo* NAD^+^ biosynthesis is inactive, and NAD^+^ pools are fueled by the salvage pathways and by the Preiss-Handler pathway. Instead, QPRT functions as an apoptosis inhibitor by interacting with caspase 3 and preventing its activation thereby conferring NSCLC resistance to apoptosis and enabling them to thrive through the stresses of tumor progression **(2)**.

Our findings also establish that NSCLC cells rely on the salvage and Preiss-Handler pathways to meet their NAD^+^ demands, with negligible contribution from the *de novo* pathway. These results are consistent with recent studies, which showed that neuroendocrine tumors and other cancers primarily depend on NAD⁺ salvage and Preiss-Handler pathways for maintaining their cellular NAD⁺ pools (6,38,61). Moreover, while the kynurenine pathway has been implicated in cancer immune evasion and metabolic adaptation (25,62), we found that QPRT’s role in NSCLC survival is independent of its function in Trp catabolism. QA accumulation or Trp restriction had no impact on cell viability, and QPRT knockdown did not affect the expression of upstream enzymes in the pathway. These findings argue against a model where toxic metabolite buildup or disrupted Trp flux drives cell death, further strengthening the conclusion that QPRT’s pro-survival function lies outside its metabolic network. Therefore, therapeutic strategies that focus solely on specific metabolic pathways may overlook critical non-metabolic survival functions of components of these pathways, such as the one we report for QPRT.

Importantly, QPRT expression positively correlates with NSCLC progression in both genetically engineered mouse models and human samples. This is in line with previous reports on breast cancer and glioma where QPRT expression was positively associated with tumor grades (63,64) and suggests a selective pressure to induce QPRT expression as tumors evolve, potentially as a strategy to counteract apoptosis. Consequently, we propose that in NSCLC cells, QPRT functions as a pro-survival scaffold, protecting them against intrinsic apoptotic signals and thereby enabling progression into higher-grade NSCLC.

These findings raise important mechanistic and translational questions. Our data showed that catalytically inactive mutants still bind caspase 3, but the interaction domains are unknown. Additional studies focused on high-resolution structural methodologies, such as X-ray crystallography or cryo-EM, can inform drug development by defining the structural interface between QPRT and caspase 3. Additionally, whether QPRT’s non-enzymatic, apoptosis-suppressive function is conserved across cancers remains a clinically important question. Our work has focused on NSCLC, but QPRT is expressed in multiple tumor types and has been linked to stress adaptation and survival in glioblastoma and breast cancer (63,64). Investigating QPRT expression, localization, and caspase 3 binding in the context of various cancer types could lead to broader therapeutic implications.

This work also highlights that metabolic enzymes can possess non-enzymatic moonlighting roles that critically influence cancer cell fate. As with other dual-function enzymes (e.g., PKM2, IDH1) (38), QPRT’s non-catalytic activity appears to be essential for tumor progression. Targeting such protein–protein interactions rather than enzymatic active sites may provide a new direction for drug development, especially for cases where catalytic inhibition has yielded limited therapeutic success. Moreover, our work suggests a context-dependent switch that regulates whether QPRT engages in its metabolic or apoptotic roles. Further studies are needed to identify the factors that regulate this switch, including post-translational modifications, protein partners, and subcellular localization cues. QPRT may serve as a regulatory node whose function switches in response to environmental or oncogenic stress, potentially creating therapeutic liabilities.

In summary, our data established that QPRT in the context of NSCLC does not only act as an enzyme in NAD⁺ biosynthesis but also as a critical apoptotic regulator that promotes NSCLC cell survival independently of NAD⁺ metabolism. This newly identified dual functionality positions QPRT as a unique therapeutic target to selectively sensitize NSCLC cells to apoptosis, a promising avenue for improving treatment outcomes in NSCLC.

## Acknowledgements

We thank members of the Gomes and DeNicola Laboratories for their helpful feedback and Skylar Robbins for technical assistance. We would also like to thank the Moffitt Cancer Center/USF Comparative Medicine Program for animal care, and staff members of the Flow Cytometry, Proteomics and Metabolomics, Chemical Biology as well as the Analytical Microscopy core facilities at the H. Lee Moffitt Cancer Center & Research Institute, an NCI designated Comprehensive Cancer Center (P30-CA076292). This work was directly supported by a Phi Beta Psi Sorority Award to APG and a joint Florida Health Department Bankhead-Coley Research Program Award (21B06) to APG, GMD and JSG. APG and the Gomes Laboratory are also supported by a New Innovator Award from OD/NIH to APG (DP2AG0776980), an American Cancer Society Research Scholar Award (RSG-22-164-01-MM), the NIA (R21AG083720), the NCI (R01CA279023), the Florida Health Department Bankhead-Coley Research Program (24B03), METAvivor, and the Florida Breast Cancer Research Foundation. SD was supported by a Miles for Moffitt Postdoctoral Fellowship as well as by a Postdoctoral Fellowship from the American Cancer Society (PF-24-1151991-01-MM) and is currently supported by a K99/R00 award from the NCI (K99CA304508). AAN was supported by NIH R35 GM143004 to JMB. Schematic figures were created using BioRender.com.

## Author Contributions

Conceptualization: APG, GMD

Methodology: HK, AAN, SD, DI

Investigation: HK, DI, AAN, SD, FL, DR, JL, JDS, NPW

Formal Analysis: HK, JL, DI, NPW, TD

Visualization: HK, AAN, DI, JL, SD, DR, NPW

Resources: APG, GMD, JSG, ERF, JMB

Funding Acquisition: APG, GMD, JSG

Project administration: APG, GMD, DI

Supervision: APG, GMD, JSG, ERF, JMB

Writing – original draft: HK, DI, APG

Writing – review & editing: HK, DI, APG, GMD, SD, JL, AAN, JMB

## Notes

### Competing Interest Statement

The authors have declared no competing interest.

