## Supplementary Fig. S1 for "Quinolinic acid phosphoribosyl transferase moonlights as an apoptosis regulator to empower lung cancer progression"

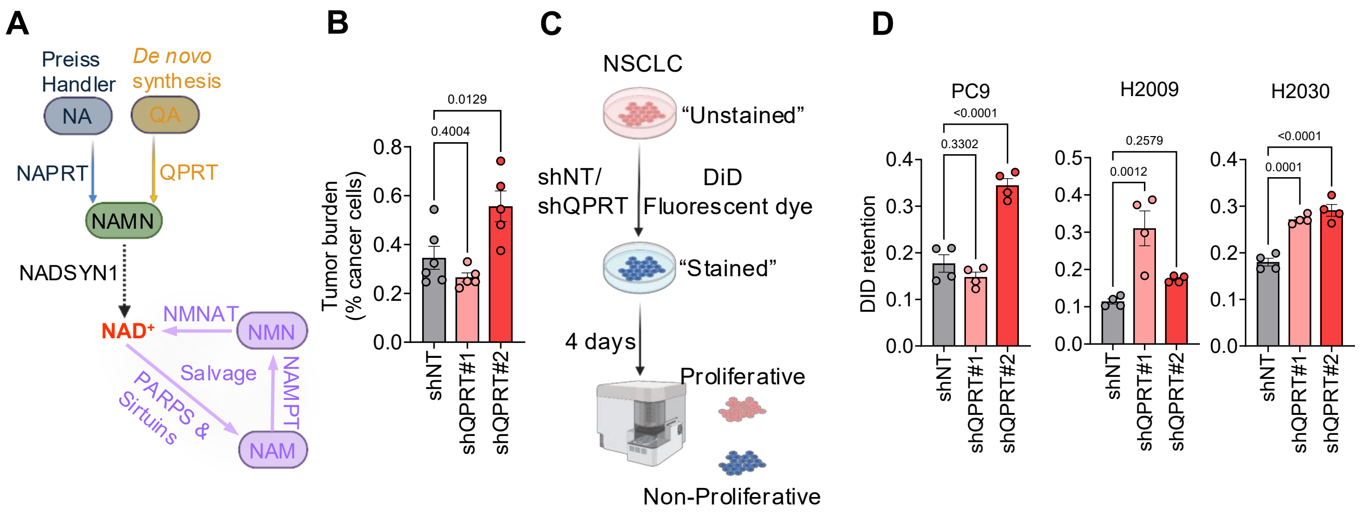


**Supplementary Figure 1. QPRT silencing suppresses NSCLC cell proliferation and induces cell death independent of cell cycle arrest. (A)** Schematic of $NAD^+$ biosynthesis pathway. **(B)** Quantification of tumor burden in H&E slides from NSG mice one week after tail vein injection with H2030 cells expressing shNT, shQPRT#1 or shQPRT#2 (shNT n = 6, shQPRT#1 n = 5 and shQPRT#2 n = 5). **(C)** Schematic of experimental design for the DiD proliferation assay. **(D)** DID proliferation assay in PC9, H2009 and H2030 upon QPRT silencing (n = 4).

Data for A and C are expressed as mean ± SEM, and statistical significance measured by ordinary one-way ANOVA test followed by Dunnett's post hoc test.
