## Supplementary Fig. S2 for "Quinolinic acid phosphoribosyl transferase moonlights as an apoptosis regulator to empower lung cancer progression"

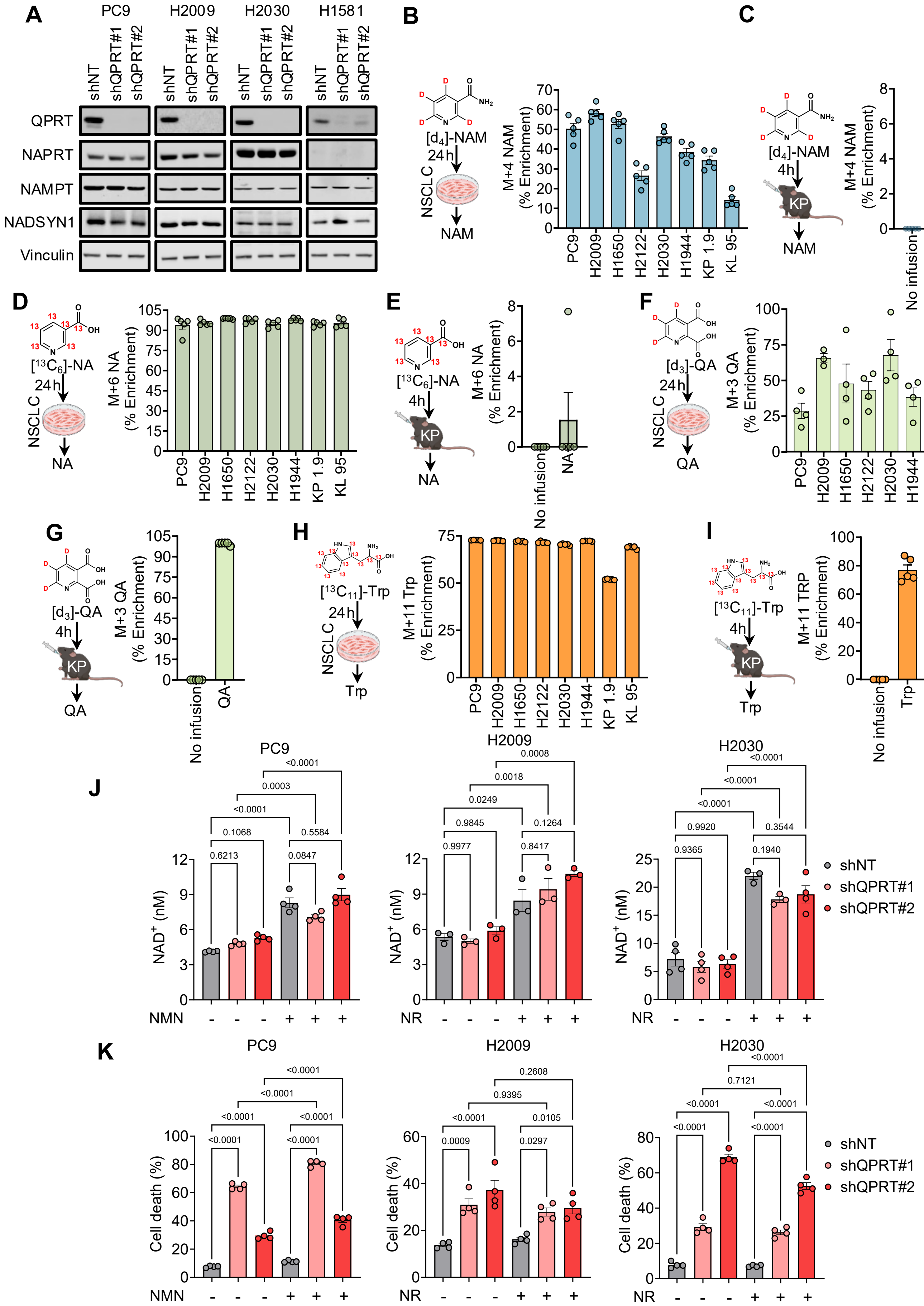
A
PC9
H2009
H2030
H1581
shNT
shQPRT#1
shQPRT#2
QPRT
NAPRT
NAMPT
NADSYN1
Vinculin
B
[d4]-NAM
24 h
NSCLC
NAM
M+4 NAM (% Enrichment)
PC9
H2009
H1650
H2122
H2030
H1944
KP 1.9
KL 95
C
[d4]-NAM
4h
KP
NAM
M+4 NAM (% Enrichment)
No infusion
NAM
D
[13C6]-NA
24 h
NSCLC
NA
M+6 NA (% Enrichment)
E
[13C6]-NA
4h
KP
NA
M+6 NA (% Enrichment)
No infusion
NA
F
[d3]-QA
24 h
NSCLC
QA
M+3 QA (% Enrichment)
G
[d3]-QA
4h
KP
QA
M+3 QA (% Enrichment)
No infusion
QA
H
[13C11]-Trp
24 h
NSCLC
Trp
M+11 Trp (% Enrichment)
I
[13C11]-Trp
4h
KP
Trp
M+11 TRP (% Enrichment)
No infusion
Trp
J
PC9
H2009
H2030
NAD+ (nM)
NMN
NR
shNT
shQPRT#1
shQPRT#2
K
PC9
H2009
H2030
Cell death (%)
NMN
NR
shNT
shQPRT#1
shQPRT#2

**Supplementary Figure 2. QPRT silencing does not alter $NAD^+$ levels or induce compensatory upregulation of $NAD^+$ biosynthetic enzymes. (A)** Immunoblot analysis showing expression of rate-limiting enzymes in the $NAD^+$ biosynthetic pathway following QPRT silencing (representative images, n = 4). **(B-I)** Stable isotope tracing *in vitro* in NSCLC cells after 24h treatment using $d_4$NAM (32 μM) for NAM **(B)**, $^{13}C_6$NA (20 μM) for NA **(D)**, $d_3$QA (20 μM) for QA **(F)**, and $^{13}C_{11}$Trp (80 μM) for Trp **(H)**, (n = 5) and *in vivo* in KP tumors with no infusion or after 4h infusion using $d_4$NAM (4 mM) for NAM **(C)**, $^{13}C_6$NA (0.2 mM) for NA **(E)**, $d_3$QA (4 mM) for QA, and **(G)**, $^{13}C_{11}$Trp (100 mM) for Trp **(I)**, (n = 5). **(J, K)** Quantification of $NAD^+$ (n = 4 for PC9, n = 3 for H2009, and n = 3-4 for H2030 ) **(J)** and cell death using PI staining for H2009 and H2030, and SYTOX Green for PC9 (n = 4 for all) **(K)** following QPRT silencing in PC9, H2009 and H2030 cells pretreated with NAD precursors, NMN (1 mM) or NR (1 mM).

Data are represented as mean ± SEM for B-J, and statistical significance measured by one-way ANOVA followed by Dunnett's post hoc test for panels J and K.
