## Supplementary Fig. S3 for "Quinolinic acid phosphoribosyl transferase moonlights as an apoptosis regulator to empower lung cancer progression"

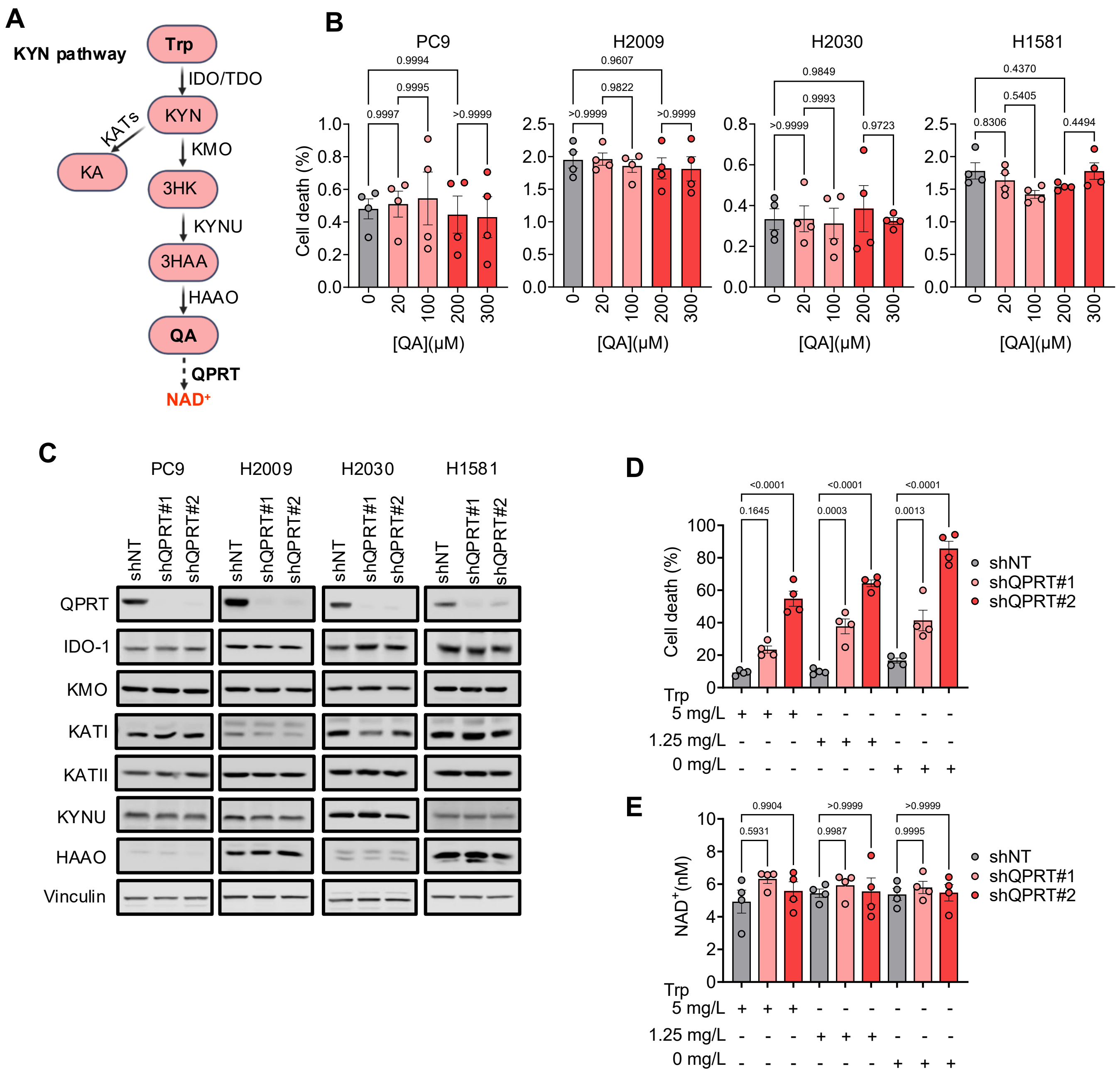


**Supplementary Figure 3. QPRT silencing does not induce cell death via kynurenine pathway metabolite accumulation or flux perturbations. (A)** Schematics of kynurenine (KYN) pathway. **(B)** Cell viability analysis with PI staining in NSCLC panel after exogenous QA treatment at varying concentrations (n = 4). **(C)** Immunoblot analysis of kynurenine pathway enzymes in NSCLC panel upon QPRT silencing (n = 3). **(D, E)** Quantification of cell death **(D)** and $NAD^+$ levels **(E)** in H2030 cells with QPRT knockdown after restricting extracellular tryptophan to final concentrations of 0 mg/L and 1.25 mg/L compared to normal levels of 5 mg/L in standard RPMI 1640 media (n = 4).

Data are represented as mean ± SEM with statistical significance measured by one-way ANOVA followed by Tukey’s post hoc test for B, D and E.
