## Supplementary Fig. S4 for "Quinolinic acid phosphoribosyl transferase moonlights as an apoptosis regulator to empower lung cancer progression"

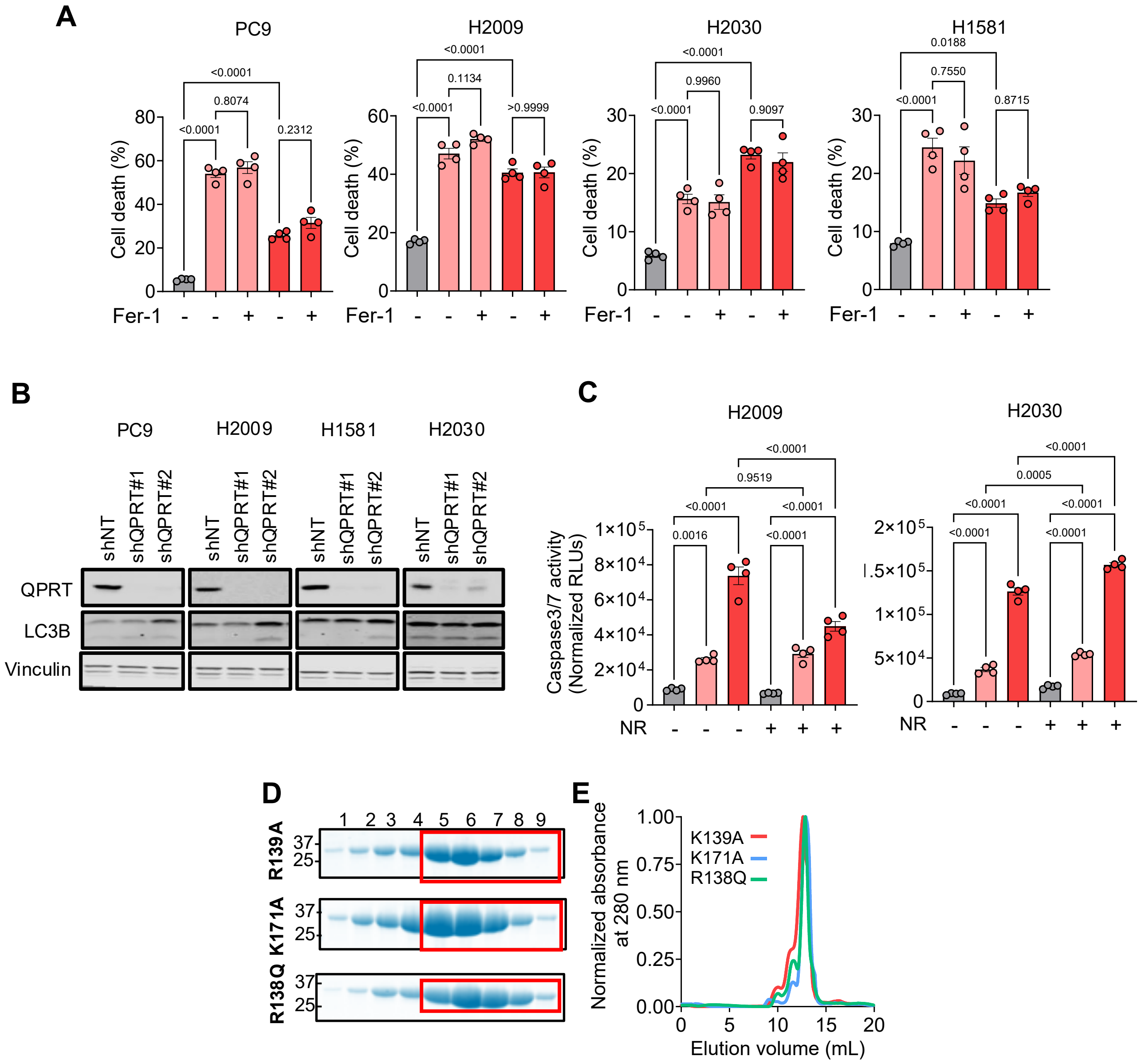


**Supplementary Figure 4. QPRT silencing does not induce ferroptosis or autophagy, but drives apoptosis independently of NAD⁺ metabolism. (A)** Cell viability analysis after treatment with ferrostatin-1 (Fer-1) in using PI staining for PC9, H2009 and H2030, and SYTOX Green for H1581 cells with QPRT silenced (n = 4). **(B)** Immunoblot analysis of LC3B levels in NSCLC cells after QPRT silencing (n = 3). **(C)** Caspase-3/7 activity levels in NR-pretreated H2009 and H2030 cells upon QPRT silencing (n = 4). **(D)** Validation of purification of K139A, K171A and R138Q mutant QPRT proteins expressed in BL21 (DE3) E. coli cells by SDS–PAGE Stain-Free detection. **(E)** Size-exclusion chromatography (SEC) profiles of the QPRT mutants. The monodisperse peaks eluting between 10 and 15 mL indicate high purity and homogeneity.

Data are represented as mean ± SEM with statistical significance measured by one-way ANOVA followed by Tukey's post hoc test for A and C.
