## Supplementary Table 1 for "Quinolinic acid phosphoribosyl transferase moonlights as an apoptosis regulator to empower lung cancer progression"

**Table S1. Cell cycle analysis of NSCLC cells upon QPRT silencing.** Data are expressed as mean ± SEM (n = 4). Statistical significance was measured by two-way ANOVA test. For each cell line and cell cycle category, p values between shNT and each shQPRT condition are displayed in parentheses.

| | CONDITION | G0/G1 | S | G2M |
|---|---|---|---|---|
| **PC9** | shNT | 71.03 ± 1.06 | 11.35 ± 0.88 | 14.45 ± 0.52 |
| | shQPRT#1 | 67.75 ± 1.23 (p = 0.0350) | 15.25 ± 1.12 (p = 0.0103) | 14.68 ± 0.72 (p = 0.9826) |
| | shQPRT#2 | 67.68 ± 1.64 (p = 0.0305) | 9.61 ± 0.77 (p = 0.3602) | 19.85 ± 0.80 (p = 0.0004) |
| **H2009** | shNT | 62.83 ± 2.78 | 18.80 ± 1.50 | 17.28 ± 1.57 |
| | shQPRT#1 | 72.28 ± 1.40 (p < 0.0001) | 15.70 ± 0.53 (p = 0.1147) | 11.25 ± 0.87 (p = 0.0014) |
| | shQPRT#2 | 62.98 ± 0.53 (p = 0.9938) | 16.48 ± 0.21 (p = 0.2722) | 18.50 ± 0.23 (p = 0.6725) |
| **H2030** | shNT | 64.03 ± 1.63 | 14.70 ± 0.30 | 18.63 ± 1.25 |
| | shQPRT#1 | 66.35 ± 0.97 (p = 0.2220) | 11.97 ± 0.89 (p = 0.1291) | 19.68 ± 0.70 (p = 0.7271) |
| | shQPRT#2 | 49.05 ± 1.75 (p < 0.0001) | 17.95 ± 0.75 (p = 0.0596) | 26.15 ± 0.80 (P < 0.0001) |
